# Endosidin20 targets cellulose synthase catalytic domain to inhibit cellulose biosynthesis

**DOI:** 10.1101/2020.02.16.946244

**Authors:** Lei Huang, Xiaohui Li, Weiwei Zhang, Nolan Ung, Nana Liu, Xianglin Yin, Yong Li, Robert E. Mcewan, Brian Dilkes, Mingji Dai, Glenn R. Hicks, Natasha V. Raikhel, Christopher J. Staiger, Chunhua Zhang

## Abstract

Cellulose is synthesized by rosette structured cellulose synthase (CESA) complexes (CSCs), each of which is composed of multiple units of CESAs in three different isoforms. CSCs rely on vesicle trafficking for delivery to the plasma membrane where they catalyze cellulose synthesis. Although the rosette structured CSCs were observed decades ago, it remains unclear what amino acids in plant CESA that directly participate in cellulose catalytic synthesis. It is also not clear how the catalytic activity of CSCs influences their efficient transport at the subcellular level. Here we report characterization of the small molecule Endosidin20 (ES20) and present evidence that it represents a new CESA inhibitor. We show data from chemical genetic analyses, biochemical assays, structural modeling, and molecular docking to support our conclusion that ES20 targets the catalytic site of Arabidopsis CESA6. Further, chemical genetic analysis reveals important amino acids that potentially form the catalytic site of plant CESA6. Using high spatiotemporal resolution live-cell imaging, we found that inhibition of CSC catalytic activity by inhibitor treatment, or by creating missense mutation at amino acids in the predicted catalytic site, causes reduced efficiency in CSC transport to the plasma membrane. Our results show that the catalytic activity of plant CSCs is integrated with subcellular trafficking dynamics.

**One sentence summary:** Endosidin20 targets cellulose synthase at the catalytic site to inhibit cellulose synthesis and the inhibition of catalytic activity reduces cellulose synthase complex delivery to the plasma membrane.

## INTRODUCTION

Cellulose is a polymer of β-1,4-D-glucose that serves as an essential cell wall component for anisotropic growth of plant cells. Cellulose is synthesized at the plasma membrane (PM) by a cellulose synthase complex (CSC) comprising a rosette of subunits in a hexagonal array with 25-nm diameter that can be observed in many different plant cell types (Mueller et al., 1976; Giddings et al., 1980; Mueller and Brown, 1980). Each CSC is predicted to contain at least 18 monomeric cellulose synthases (CESAs) of three different isoforms in a 1:1:1 molar ratio (Pear et al., 1996; Arioli et al., 1998; Doblin et al., 2002; Persson et al., 2007; Fernandes et al., 2011; Newman et al., 2013; Gonneau et al., 2014; Hill et al., 2014). Freeze-fracture electron microscopy analysis reveals that CSCs are localized to the Golgi, Golgi-derived vesicles and the PM, indicating CSCs are transported through the vesicle trafficking pathway (Haigler and Brown, 1986). Live cell imaging of CSCs using functional fluorescence-tagged CESA provides further compelling evidence that CSCs are localized to the Golgi/trans-Golgi network (TGN), small CESA-containing compartments (SmaCCs)/microtubule-associated cellulose synthase compartments (MASCs), and PM (Paredez et al., 2006; Crowell et al., 2009; Gutierrez et al., 2009; Zhang et al., 2019). PM-localized CSCs show bidirectional motility and the velocity is related to cellulose polymerization process (Paredez et al., 2006; Crowell et al., 2009; Gutierrez et al., 2009; Fujita et al., 2013).

Precise CSC delivery to the PM requires the coordinated function of multiple cellular machineries including microtubules, actin, and general exocytosis machinery (Paredez et al., 2006; Crowell et al., 2009; Farquharson, 2009; Gutierrez et al., 2009; Sampathkumar et al., 2013; Bashline et al., 2014; McFarlane et al., 2014; Lei et al., 2015; Luo et al., 2015; Zhu et al., 2018; Zhang et al., 2019). CSCs in the Golgi and SmaCCs require actin cytoskeleton for long and short distance cellular transport (Crowell et al., 2009; Gutierrez et al., 2009; Sampathkumar et al., 2013). SmaCCs that are close to the PM interact with microtubules and CELLULOSE SYNTHASE INTEACTIVE 1 (CSI1) for targeted delivery to the PM (Farquharson, 2009; Gutierrez et al., 2009; Li et al., 2012). CSCs also require the conserved exocyst complex, PATROL1, actin, and myosin XI for tethering and fusion to the PM (Sampathkumar et al., 2013; Zhu et al., 2018; Zhang et al., 2019). There are also new cellular components, such as STELLO and SHOU4, that regulate CSC delivery to the PM through molecular mechanisms that require further investigation (Zhang et al., 2016b; Polko et al., 2018). High resolution structural analysis has defined key amino acids in the catalytic site of *Rhodobacter sphaeroidesin* cellulose synthase (RsBcsA) (Morgan et al., 2013; Omadjela et al., 2013; Morgan et al., 2014; Morgan et al., 2016). However, plant CESAs contain plant-conserved sequence (P-CR) and class-specific region (CSR) in the cytoplasmic domain that are not present in bacterial CesA (Pear et al., 1996; Vergara and Carpita, 2001). The presence of P-CR and CSR indicates that the plant CESA catalytic site may have different amino acid composition compared with the bacterial CesA catalytic site. So far, high resolution structures for CSCs and individual CESAs are not available to define amino acids that participate in cellulose catalytic synthesis. It also remains inconclusive whether the catalytic site contains information that influences CSC delivery to the PM.

Small molecule inhibitors of CESA, for example isoxaben and dichlorobenzonitrile (DCB), have proven useful in understanding the molecular functions and dynamics of CSCs (Montezinos and Delmer, 1980; Heim et al., 1989; Scheible et al., 2001; Desprez et al., 2002; DeBolt et al., 2007b; Brabham et al., 2014; Worden et al., 2015; Tateno et al., 2016; Tran et al., 2018). However, there are limitations to using these small molecules because the mechanisms of how these inhibitors target CESAs directly are not known. Here, we report the characterization of a new small molecule, Endosidin20 (ES20), that inhibits cellulose synthesis by directly targeting Arabidopsis CESA6. We use chemical genetic analyses, structural modeling, molecular docking and biochemical assays to show that ES20 targets CESA6 at the catalytic site that requires proper functions of at least 18 amino acids in the cytoplasmic domain. Further, we analyzed the cellular localization and trafficking dynamics of CSCs in ES20-treated seedlings and CSCs that contain missense mutations at the catalytic site of CESA6. We found that the trafficking dynamics of CSC is altered after ES20 treatment or when the catalytic site is mutated, indicating that the catalytic site of CSC contains information that affects their efficient subcellular transport.

## RESULTS

### Endosidin20 inhibits cellulose synthesis

Endosidin20 (ES20) (Figure 1A) was initially found to inhibit pollen tube growth without interfering with the trafficking of two types of cargo proteins, PIN2 auxin transporter and BRI1 brassinosteroid receptor, that are delivered to and maintained at the PM (Figure 1B) (Drakakaki et al., 2011). ES20 also did not disrupt the localization of two other plasma membrane proteins, ROP6 and PIP2a, or general marker proteins for endoplasmic reticulum (ER) (GFP-HDEL), Golgi (YFP-Got1p), and trans-Golgi Network/Early Endosomes (TGN/EEs) (VHA-a1-GFP) (Figure 1B). We found that ES20 induced plant growth phenotypes similar to cellulose synthase mutants or wildtype plants grown in the presence of cellulose synthase inhibitor (Arioli et al., 1998; DeBolt et al., 2007a). When grown in the presence of ES20, dark-grown hypocotyls of Arabidopsis wildtype (ecotype Col-0) seedlings became shorter and wider in an ES20 dose-dependent manner (Figure 2A-2E). The epidermal cells of hypocotyls grown in the presence of ES20 were swollen (Figure 2F-2H). In addition to these hypocotyl phenotypes, ES20 inhibited root growth in a dose-dependent manner (Figure 2I, 2J). When treated with ES20 overnight, the root tip region of wildtype plants was swollen and root elongation was significantly inhibited compared to mock-treated roots (Figure 2K-2N). Similar to hypocotyl epidermal cells, epidermal cells from the root elongation zone were markedly swollen after ES20 treatment, which was reflected by a significantly decreased cell length and a significantly increased cell width (Figure 2O-2Q). Swollen cells and organs in plants are often caused by direct or indirect disruption of cell wall biosynthesis or organization such as those caused by CESA-deficient mutants or in wildtype plants treated with inhibitors of cellulose synthesis or microtubule organization (Baskin et al., 1994; Arioli et al., 1998; Fagard et al., 2000; Burn et al., 2002; Desprez et al., 2002; Daras et al., 2009). The observation of reduced cell elongation and inhibition of anisotropic growth motivated us to test the effects of ES20 on cellulose synthesis. We found that ES20 reduced the crystalline cellulose content of both dark-grown hypocotyls and light-grown roots of wildtype seedlings in a dose-dependent manner (Figure 2R, 2S). These results indicate that ES20 could inhibit plant growth by affecting cellulose biosynthesis. We also found that lignin and callose were accumulated at higher levels compared with control seedlings grown in the presence of DMSO (Figure 2T-U), similar to previous reports for plants with cellulose synthesis deficiency caused by mutation or inhibitors (Desprez et al., 2002; Cano-Delgado et al., 2003; Sampathkumar et al., 2013).

**Figure 1.**
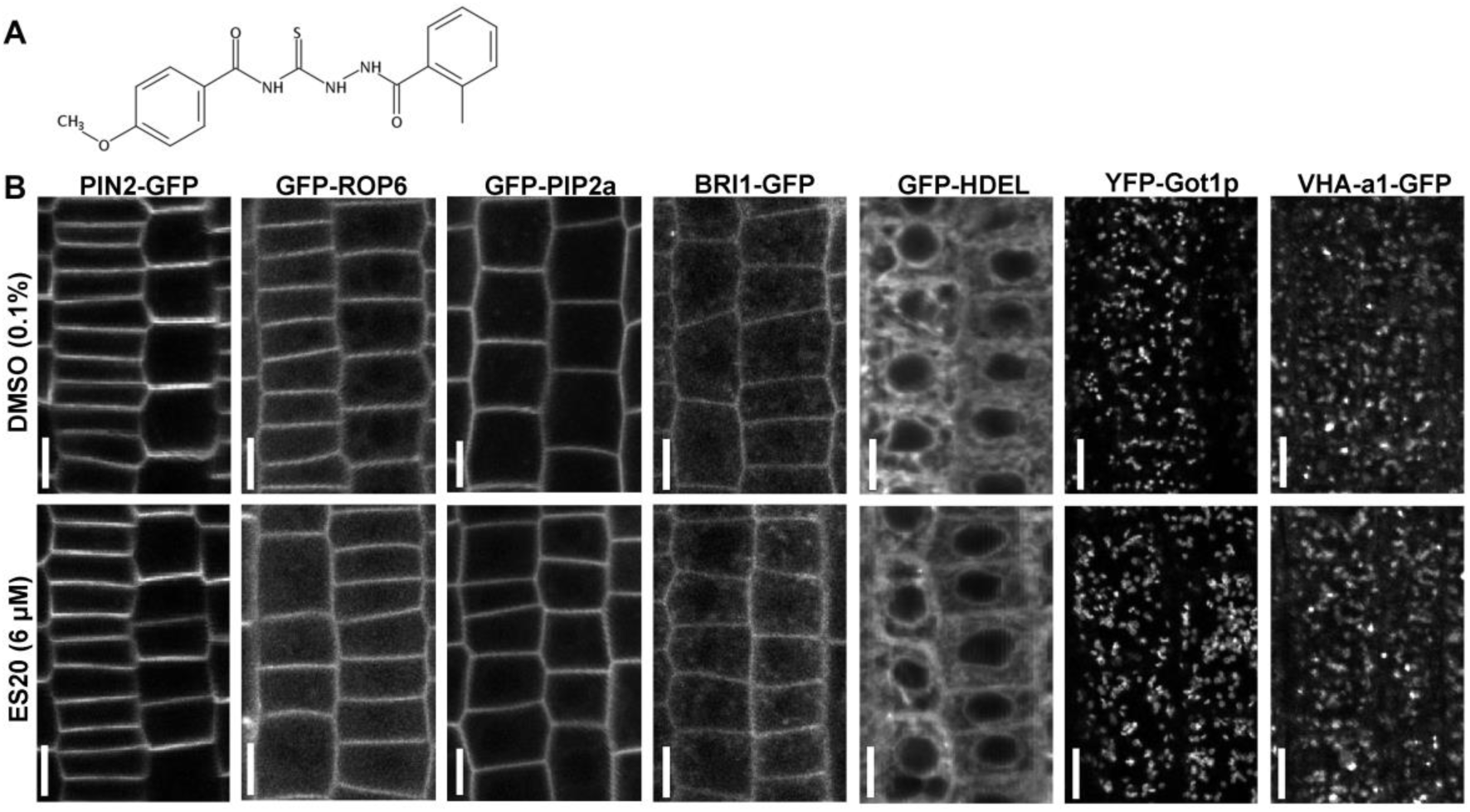
ES20 does not disrupt the general endomembrane system of plants. **A**, Molecular structure of ES20. **B**, Representative images of cellular localization of different organelle marker proteins in 5-day-old transgenic plants after treatment with 0.1% DMSO or 6 μM ES20 for 2 h in root epidermal cells from the elongation zone. PIN2-GFP, GFP-ROP6, GFP-PIP2a, and BRI1-GFP were used as PM marker proteins. GFP-HDEL was used as an ER marker protein. YFP-Got1p (Golgi transporter 1 protein) was used as a Golgi marker protein. VHA-a1-GFP (v-type proton ATPase subunit a1) was used as a TGN/EE marker protein. Scale bars: 10 μm.

**Figure 2.**
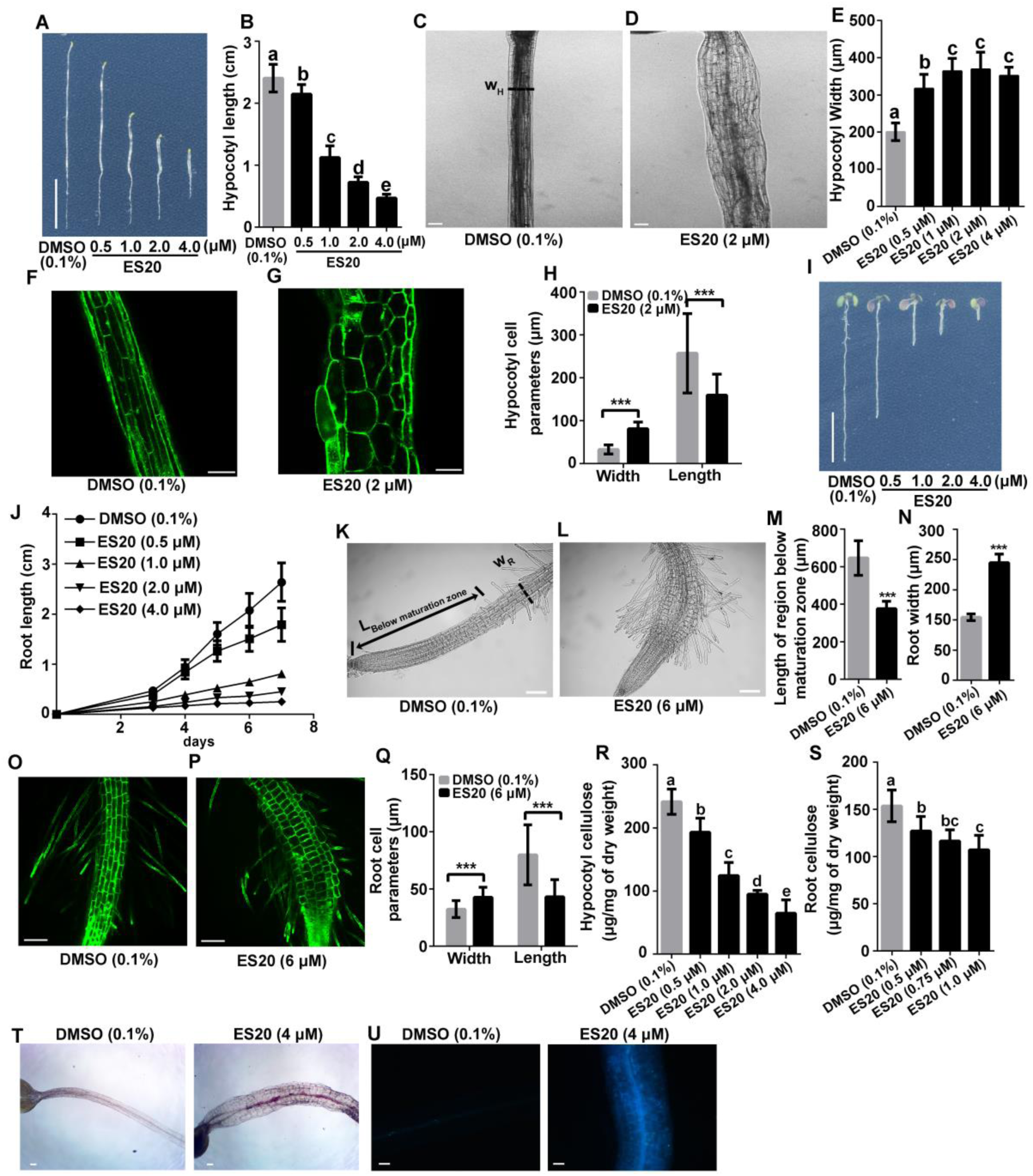
ES20 is a cellulose synthesis inhibitor. **A** and **B**, ES20 inhibits *Arabidopsis* hypocotyl growth in the dark in a dose-dependent manner. **C** to **E**, ES20 caused dark-grown hypocotyls to swell. W_H_ in **C**, the region where hypocotyl width was measured. **F** to **H,** ES20 inhibited the anisotropic growth of hypocotyl epidermal cells. **I** and **J**, ES20 inhibited *Arabidopsis* root growth in a dose-dependent manner. **K** to **N**, ES20 inhibited root growth and caused roots to swell. The tip region of 5-day-old wildtype roots treated with 0.1% DMSO (**K**) or 6 μM ES20 (**L**) for 12 hours. L_Below maturation zone_ and W_R_ in **K**, the regions where root length below maturation zone and root width were measured for **M** and **N**, respectively. **O** to **Q**, ES20 inhibited root growth and caused root cells to swell. **R** and **S**, ES20 reduced crystalline cellulose content in cell walls of dark-grown hypocotyls (**R**) and light-grown roots (**S**) in a dose-dependent manner. **T**. ES20 treatment causes accumulation of lignin. Representative images of 3 days old Col-0 seedlings grown on 1/2 MS supplemented with DMSO or 4 μM ES20 in dark stained with Phloroglucinol. **U**, ES20 treatment causes accumulation of callose. Representative images of 3 days old Col-0 seedlings grown on ½ MS supplemented with DMSO or 4 μM ES20 in dark stained with aniline blue. The letters in **B**, **E**, **R**, and **S** indicate statistically significant differences determined by one-way ANOVA tests followed by Tukey’s multiple comparison tests in different samples. Different letters indicate significant differences between groups (p < 0.05). *** in **H, M, N** and **Q**, p < 0.001 by two-tailed student’s *t* test. Error bars: mean ± SD, with *n* = 20 for **B** and **E**, *n* = 15 for **H**, **J**, **M, N** and **Q**, and *n* = 9 for **R** and **S**. Scale bars in **A** and **I** are 1.0 cm; Scale bars in other panels: 100 μm.

### Mutations at the CESAs cause reduced sensitivity to ES20

To identify the cellular and molecular pathways that are targeted by ES20, we performed a chemical genetic screen for mutants with reduced sensitivity to this growth inhibitor. We screened about 500,000 M2 seedlings from ethyl methanesulfonate (EMS) mutagenized populations on 5 μM ES20 and identified seedlings with longer roots than control plants that were not mutagenized. Selected M2 seedlings were transferred to soil to produce the M3 generation. After re-testing the seeds from the M3 generation, we confirmed a total of 45 individual lines that had reduced sensitivity to ES20 for growth. We refer to these alleles as *ES20 RESISTANT* or *es20r* genes. After high-throughput whole genome sequencing of pooled seedlings from mapping populations, we found some of these individual lines carried the same mutation. We obtained 15 different mutant alleles carrying either C-T or G-A mutations in *CESA6* and these mutants were named *es20r1*–*es20r15*. Due to the number of mutants we found in *CESA6* and associated resources available in *Arabidopsis*, we decided to use these mutants to further analyze the mode of action of ES20 and the importance of mutated amino acids for CESA6 function. We backcrossed the mutants and consistently detected reduced sensitivity of these mutants to ES20 for light-grown root and dark-grown hypocotyl growth (Figure 3A, 3B; Supplemental Figure, Figure S1; Supplemental Figure, Figure S2). By further analyzing the *es20r*/*cesa6* mutations, we found that 12 of them led to missense mutations in amino acids located at the predicted central cytoplasmic domain, two caused missense mutations in amino acids located at the predicted transmembrane domains, and one was mutated in the cytosolic domain close to the fifth predicted transmembrane domain (Figure 3C). To confirm that the mutations in *CESA6* caused reduced sensitivity to ES20, we performed genetic complementation experiments. We cloned the genomic content of *CESA6* and inserted a YFP tag at the 5’ end of the coding region so that we could generate CESA6 with an N-terminal YFP fusion. We used this wildtype YFP-CESA6 construct as a template and performed site-directed mutagenesis to create ten additional clones of YFP-CESA6, each carrying one of the mutations that were found in our EMS mutants. We transformed each wildtype and mutated YFP-CESA6 construct into a loss-of-function *cesa6* allele, *prc1-1* (Fagard et al., 2000). We selected independent transformants that contain single insertion for each construct based on the segregation ratio of selection maker in T2 plants. After analyzing the homozygous transformants with single CESA6 construct insertion, we found that the wildtype CESA6 construct could rescue the growth phenotypes of *prc1-1* in roots and hypocotyls and the mutated CESA6 constructs could fully or partially rescue the growth phenotype of *prc1-1*, depending on the mutation (Figure 3D, 3E; Supplemental Figure, Figure S3). The transgenic lines carrying any of the ten mutated CESA6 constructs exhibited reduced sensitivity to ES20 for both root and hypocotyl growth, but not the wildtype CESA6 construct (Figure 3D, 3E, Supplemental Figure, Figure S3). These genetic complementation experiments further confirmed that the missense mutations in *CESA6* are sufficient to cause reduced sensitivity to ES20.

**Figure 3.**
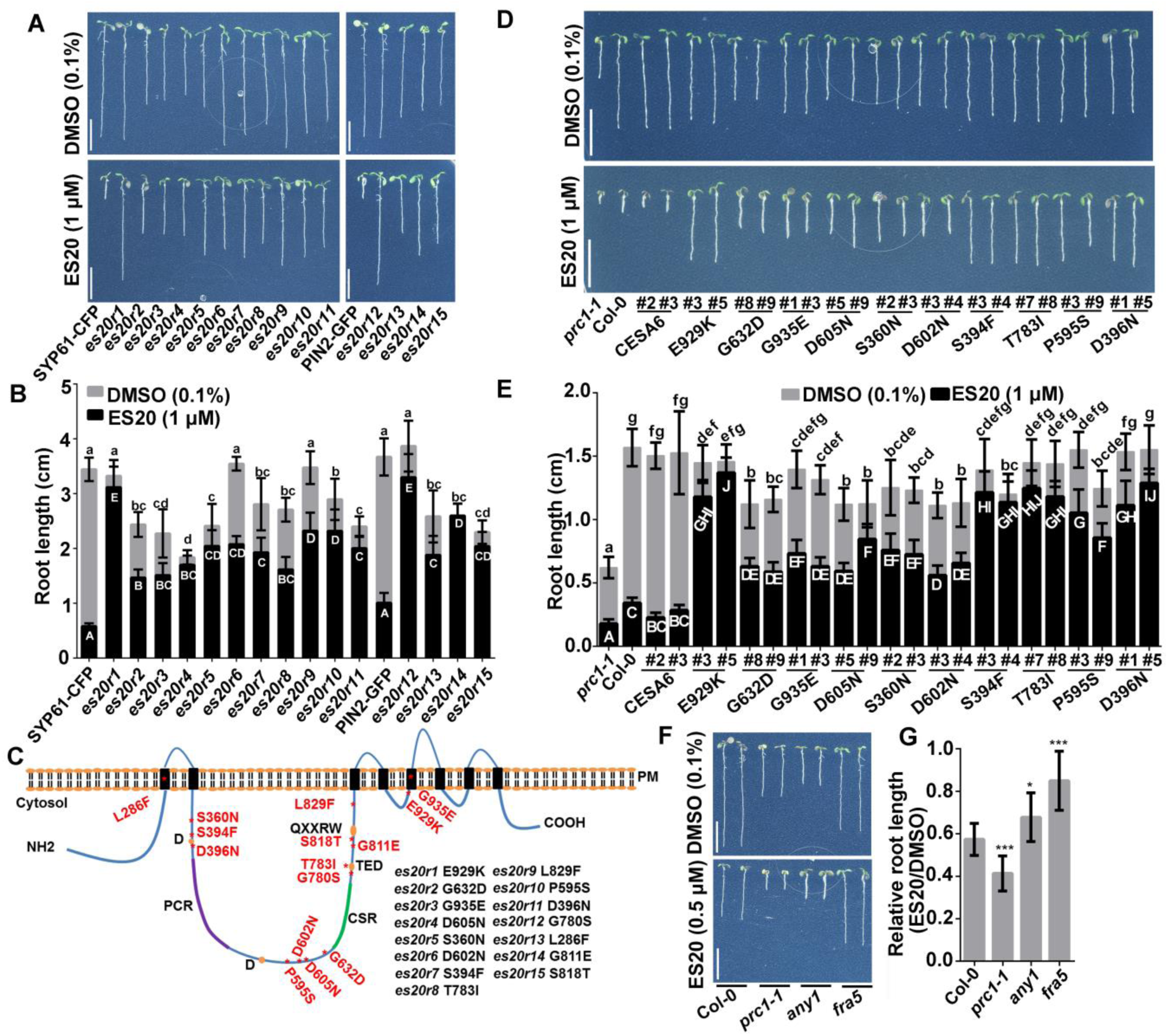
A novel collection of *cesa6* mutants have reduced sensitivity to ES20. **A**, Representative 7-day-old seedlings of wildtype expressing SYP61-CFP or PIN2-GFP, and *cesa6* mutant lines (*es20rs*) grown on media supplemented with 0.1% DMSO (top) or 1 μM ES20 (bottom). **B**, Quantification of root length of seedlings from SYP61-CFP, PIN2-GFP, and *es20r* lines grown on media supplemented with 0.1% DMSO (gray) or 1 μM ES20 (black). **C**, The diagram shows the predicted topology of CESA6 and the location of the mutated amino acid in each *es20r* allele. **D**, Genetic complementation of *prc1-1/cesa6* growth defects and sensitivity to ES20 by mutated CESA6 constructs. Ten constructs that we have tested rescued the root growth defect of *prc1-1* to different extents in the absence of ES20 and led to reduced sensitivity to ES20 in transgenic plants. **E**, Quantification on the root length of genetic complementation lines of *prc1-1/cesa6* by mutated CESA6 constructs and their sensitivity to ES20 in growth. Ten constructs that we have tested rescued the root growth defect of *prc1-1* to different extents and led to reduced sensitivity to ES20 in transgenic plants. **F** and **G**, Mutations in other CESAs also led to reduced sensitivity to ES20. The letters in **B** and **E** indicate statistically significant differences determined by one-way ANOVA tests followed by Tukey’s multiple comparison tests in different samples. Different letters indicate significant differences between groups (p < 0.05). In **B** and **E**, lower- and upper-case letters represent ANOVA analysis of plants grown on media with DMSO and ES20, respectively. Scale bars: 1 cm. Error bars represent mean ± SD, *n* = 12 in **B,** *n* = 14 in **E** and **G**. * indicates p < 0.05 and *** indicates p < 0.001 by two-tailed student’s *t* test in comparison with Col-0.

After we cloned the mutant genes, we found all these amino acids that were mutated in *es20r* mutants are conserved in *Arabidopsis* CESAs and six are conserved when compared to RsBcsA (Supplemental Figure, Figure S4). In addition, we found that the loss-of-function *cesa6* allele, *prc1-1*, was more sensitive to ES20 treatment when compared with the wildtype (Figure 3F, 3G, Supplemental Figure, Figure S5). Increased sensitivity of *prc1-1* to ES20 indicates that ES20 might target CESA2 and CESA5 that function redundantly with CESA6 (Desprez et al., 2007), or even other CESAs as well. We hypothesized that missense mutations at conserved amino acids in other CESAs in *Arabidopsis* might lead to reduced sensitivity to ES20 as well. Previously, *fra5* was reported to carry a missense mutation at P557 of *Arabidopsis* CESA7 (CESA7^P557T^) (Zhong et al., 2003), which is homologous to the conserved P595 in CESA6 (Supplemental Figure, Figure S4). Another mutant, *any1*, was reported to carry a missense mutation at D604 of *Arabidopsis* CESA1 (CESA1^D604N^) (Fujita et al., 2013), which is homologous to the conserved D605 in CESA6. We obtained *fra5* (CESA7^P557T^) and *any1* (CESA1^D604N^) and tested their response to ES20 in growth assays. As reported previously, both *fra5* (CESA7^P557T^) and *any1* (CESA1^D604N^) have shorter roots when compared with wildtype (Figure 3F), suggesting that they play a role in normal seedling growth, although CESA7 is thought to be mainly involved in secondary cell wall synthesis. We found that both *fra5* (CESA7^P557T^) and *any1* (CESA1^D604N^) showed inhibited growth upon ES20 treatment. However, when a lower concentration of ES20 (0.5 μM) was tested, the root growth of *fra5* (CESA7^P557T^) and *any1* (CESA1^D604N^) was inhibited by ES20 at a reduced level when compared with wildtype plants (Figure 3F, 3G; Supplemental Figure, Figure S5). Reduced sensitivity to ES20 in *CESA7* and *CESA1* mutants with alteration of amino acids homologous to our mutants in *CESA6* indicates that reduced sensitivity to ES20 is not unique to CESA6, but also occurs with other CESAs.

### ES20 targets the catalytic site of CESAs

To better understand how multiple mutations at conserved amino acids in CESAs affect plant sensitivity to ES20, we used a threading method to model the structure of Arabidopsis CESA6 central cytoplasmic domain using the solved crystal structure of RsBcsA as a guide (Morgan et al., 2013; Morgan et al., 2016). The reason to use RsBcsA structure as a guide is because it is the only cellulose synthase protein structure that has been solved so far, and the bacterial cellulose synthase shares many key amino acids with plant CESAs and the conservation has allowed cloning of the first plant CESA gene (Pear et al., 1996). Threading method has been used previously to predict plant CESA structure using bacterial sequences as a guide (Sethaphong et al., 2013; Nixon et al., 2016). The modeled structure of CESA6 catalytic domain contains multiple α-helices folded into a globular structure with a central cavity (Figure 4A). The quality of the model was evaluated with the PROCHECK program (Laskowski et al., 1993; Laskowski et al., 1996). In the Ramachandran plot, 67.2% of the residues were in the most favored regions and 23.4% of the residues were in the additional allowed regions. Ligand binding site prediction enabled by the COACH program identified UDP-glucose phosphonate as a possible ligand for the modeled structure of CESA6 (Yang et al., 2013a). We further used a molecular docking approach to predict the possible binding sites for ES20 on the modeled CESA6 central cytoplasmic domain structure. We found that ES20 and UDP-glucose phosphonate were docked to the same pocket in the central region of the modeled CESA6 cytoplasmic domain (Figure 4A, 4B): UDP-glucose phosphonate was docked to a predicted catalytic site that contains 25 amino acids within a distance of 4 Å (S360, T361, V362, D363, P364, K366, D396, L401, K537, K538, D562, C563, D564, H565, Q596, F598, V631, G632, T633, D785, S812, R820, Q823, R826, W827) and ES20 was docked to the same pocket that contains 18 amino acids within a distance of 5 Å (S360, T361, V362, P364, K537, K538, D562, D564, Q596, F598, G632, T633, D785, S812, R820, Q823, R826, W827). When we examined the position of mutated amino acids identified in our reduced sensitivity mutants, we found that most of these amino acids were either directly located at or very close to the predicted binding site for ES20 and UDP-glucose (Figure 4A, 4B). After further analysis of the docking results, we found that three amino acids, S360, D562 and Q823 were close to ES20 and hydrogen bonds could form between ES20 and these amino acids (Figure 4C). We also found that some of the mutated amino acids that led to reduced sensitivity to ES20 in *es20r* plants, such as D396 and T783, were conserved and the homologous amino acids were located at the catalytic core in RsBcsA and participated in the catalytic process (Morgan et al., 2016) (Supplemental Figure, Figure S4). The structural modeling and molecular docking data in combination with the chemical genetics results indicate that ES20 could target the catalytic site of CESAs to inhibit plant cellulose synthesis and cell growth.

**Figure 4.**
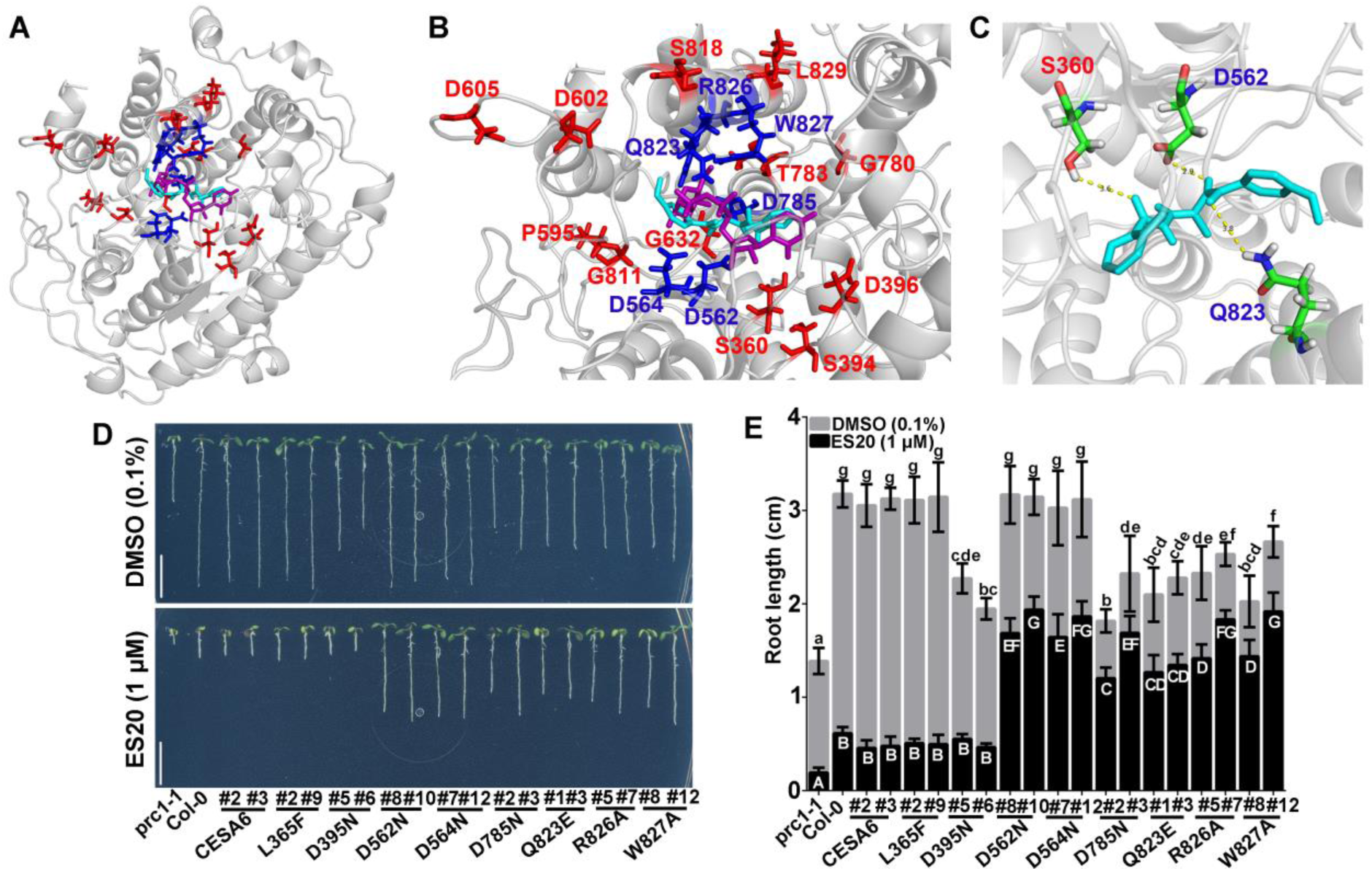
ES20 targets CESA6 at the catalytic site. **A**, The superposition of ES20 (cyan) and UDP-glucose phosphonate (magenta) on the predicted binding pocket of modeled CESA6 large cytosolic domain (amino acids 322-868). The amino acids that were mutated in *es20r* mutants (red) and the predicted amino acids (blue) that caused reduced sensitivity to ES20 when mutated were shown as sticks. **B**, Magnified view of (**A**) at the predicted binding pocket for ES20 (cyan), UDP-glucose phosphonate (magenta) and amino acids that were required for ES20 sensitivity (red and blue). **C**, The possible hydrogen bonds that were predicted to form between ES20 and S360, D562, and Q823 of CESA6. ES20 was shown as sticks and colored in cyan. **D** and **E**, Mutation of six amino acids at the predicted binding site caused reduced sensitivity to ES20. The genomic construct for CESA6^L365F^ completely rescued, whereas CESA6^D395N^ partially rescued the growth of *prc1-1,* when compared to the wildtype CESA6 construct. The transgenic plants expressing CESA6^L365F^ and CESA6^D395N^ had similar levels of sensitivity to ES20 as those expressing wildtype CESA6. The genomic constructs of CESA6^D562N^, CESA6^D564N^, CESA6^D785N^, CESA6^Q823E^, CESA6^R826A^, and CESA6^W827A^ rescued the growth of *prc1-1* to different extents and led to reduced sensitivity to ES20 in transgenic plants. The letters in **E** indicate statistically significant differences determined by one-way ANOVA tests followed by Tukey’s multiple comparison tests in different samples. Different letters indicate significant differences between groups (p < 0.05). In **E**, lower- and upper-case letters represent ANOVA analysis of plants grown on media with DMSO and ES20, respectively. Error bars: mean ± SD, with *n* = 15 in **E**. Scale bars: 1 cm.

To further validate our structural model and molecular docking data, we hypothesized that if we mutated other amino acids in the predicted binding site, the plants should have reduced sensitivity to ES20. We selected six amino acids that were located at the predicted ES20 and UDP-glucose binding site on CESA6 (Figure 4A, 4B, amino acids colored blue), including D562 and Q823 that are predicted to be important for ES20 interaction with CESA6, and created six YFP-CESA6 genomic constructs that each carried a missense mutation in one of these six amino acids. We also selected L365 and D395 that are not within 4 Å of the predicted UDP-glucose binding site nor within 5 Å of predicted ES20 binding site and created YFP-CESA6 genomic constructs that each carried a missense mutation in one of the two amino acids. We then transformed these constructs into the *prc1-1/cesa6* loss-of-function mutant and obtained single insertion transgenic lines expressing each of the mutated CESA6 containing a predicted missense mutation. In the absence of ES20, transgenic plants expressing a wildtype CESA6 construct had normal root and hypocotyl growth when compared to wildtype controls, whereas the transgenic plants expressing the mutated CESA6 had different severity of growth defects, depending on the mutation (Figure 4D, 4E; Supplemental Figure, Figure S6; Supplemental Figure, Figure S7). We analyzed YFP-CESA6 protein level in transgenic lines expressing wildtype or mutated YFP-CESA6 and found that the severity of growth defects is not corelated with the protein level (Supplemental Figure, Figure S8). In the presence of ES20, transgenic plants expressing wildtype CESA6 constructs had similar sensitivity to ES20 for root and hypocotyl growth compared to wildtype plants (Figure 4D, 4E; Supplemental Figure, Figure S6; Supplemental Figure, Figure S7). The transgenic plants expressing 6 mutated CESA6 constructs in amino acids (D562N, D564N, D785N, Q823E, R826A, W827A) within 4 Å of the predicted UDP-glucose binding site and within 5 Å of predicted ES20 binding site showed reduced sensitivity to ES20 for both root and hypocotyl growth (Figure 4D, 4E; Supplemental Figure, Figure S6; Supplemental Figure, Figure S7), suggesting that these 6 amino acids are important for the inhibitory effect of ES20. The transgenic plants expressing YFP-CESA6 carrying mutations at the two amino acids (L365F and D395N) that are not within 4 Å of predicted UDP-glucose binding site and 5 Å of predicted ES20 binding site have the same sensitivity to ES20 inhibition as wildtype YFP-CESA6, indicating these two amino acids are not essential for the inhibitory effect of ES20 (Figure 4D, 4E; Supplemental Figure, Figure S6; Supplemental Figure, Figure S7). The CESA6 protein level in different transgenic lines is not affected by ES20 treatment (Supplemental Figure S8). Together with the 12 mutant alleles that were identified through EMS mutant screening, a total of 18 alleles have been obtained that have mutations around the predicted binding site and exhibited reduced sensitivity to ES20. Transgenic lines expressing CESA6^L365F^ showed similar growth rates and similar sensitivity to ES20 in roots and hypocotyls when compared to wildtype plants (Figure 4D, 4E; Supplemental Figure, Figure S6; Supplemental Figure, Figure S7), suggesting this amino acid is not critical for plant growth or the inhibitory activity of ES20. The construct CESA6^D395N^ could partially rescue the growth of *prc1-1* and the transgenic plants had normal sensitivity to ES20 (Figure 4D, 4E; Supplemental Figure, Figure S6; Supplemental Figure, Figure S7), suggesting CESA6D395 is required for cell growth but not for the inhibitory activity of ES20. Reduced sensitivity to ES20 for plant growth caused by six mutations predicted based on the modeled structure and molecular docking provides additional evidence that ES20 targets the putative catalytic site of CESA6.

We next used biochemical assays to further test whether ES20 targets CESA6 directly. We first tried a Drug Affinity Responsive Target Stability (DARTS) assay, which detects small molecule and protein interaction by testing whether the small molecule protects the protein from degradation by proteases (Lomenick et al., 2009; Zhang et al., 2016a). Because a mixture of different types of proteases is often used against a protein or a mixture of different proteins in DARTS assay, it is a more qualitative assay than being quantitative for binding kinetics. Instead, examples from multiple protein and small molecule interactions have shown that often one or two concentrations of pronase, a mixture of different types of proteases, show significant protection of the target protein (Chin et al., 2014; Qu et al., 2016; Zhang et al., 2016a; Li et al., 2017; Kania et al., 2018; Mishev et al., 2018; Dejonghe et al., 2019; Rodriguez-Furlan et al., 2019; Zou et al., 2019). The concentrations of the ligand molecule used in DARTS assay are typically higher than the biological inhibitory concentrations (Chin et al., 2014; Qu et al., 2016; Zhang et al., 2016a; Li et al., 2017; Kania et al., 2018; Mishev et al., 2018; Dejonghe et al., 2019; Rodriguez-Furlan et al., 2019; Zou et al., 2019). We isolated total proteins from seedlings of YFP-CESA6 transgenic line and then incubated the proteins with either ES20 or DMSO control. After incubation with the small molecules, the protein was digested with pronase. We used anti-GFP antibody to detect the abundance of YFP-CESA6 after pronase digestion. We found that ES20 significantly protected YFP-CESA6 from degradation by pronase (Figure 5A, 5B). The control molecule, Ampicillin, did not protect YFP-CESA6 from degradation by pronase (Figure 5C, 5D). ES20 protection of YFP-CESA6 from degradation by proteases suggests that ES20 and YFP-CESA6 physically interact. To test whether the mutations in *es20r*s affect the interaction between CESA6 and ES20, we performed DARTS assay using total protein isolated from seedlings of YFP-CESA6^P595S^ transgenic line and ES20. We chose YFP-CESA6^P595S^ for the test because *esr20-10* (CESA6^P595S^) shows strong resistance to ES20 treatment in growth (Figure 3A-3C). We found that ES20 did not protect YFP-CEAS^P595S^ from degradation by pronase (Figure 5E, 5F), indicating P595 is important for the interaction between ES20 and CESA6.

**Figure 5.**
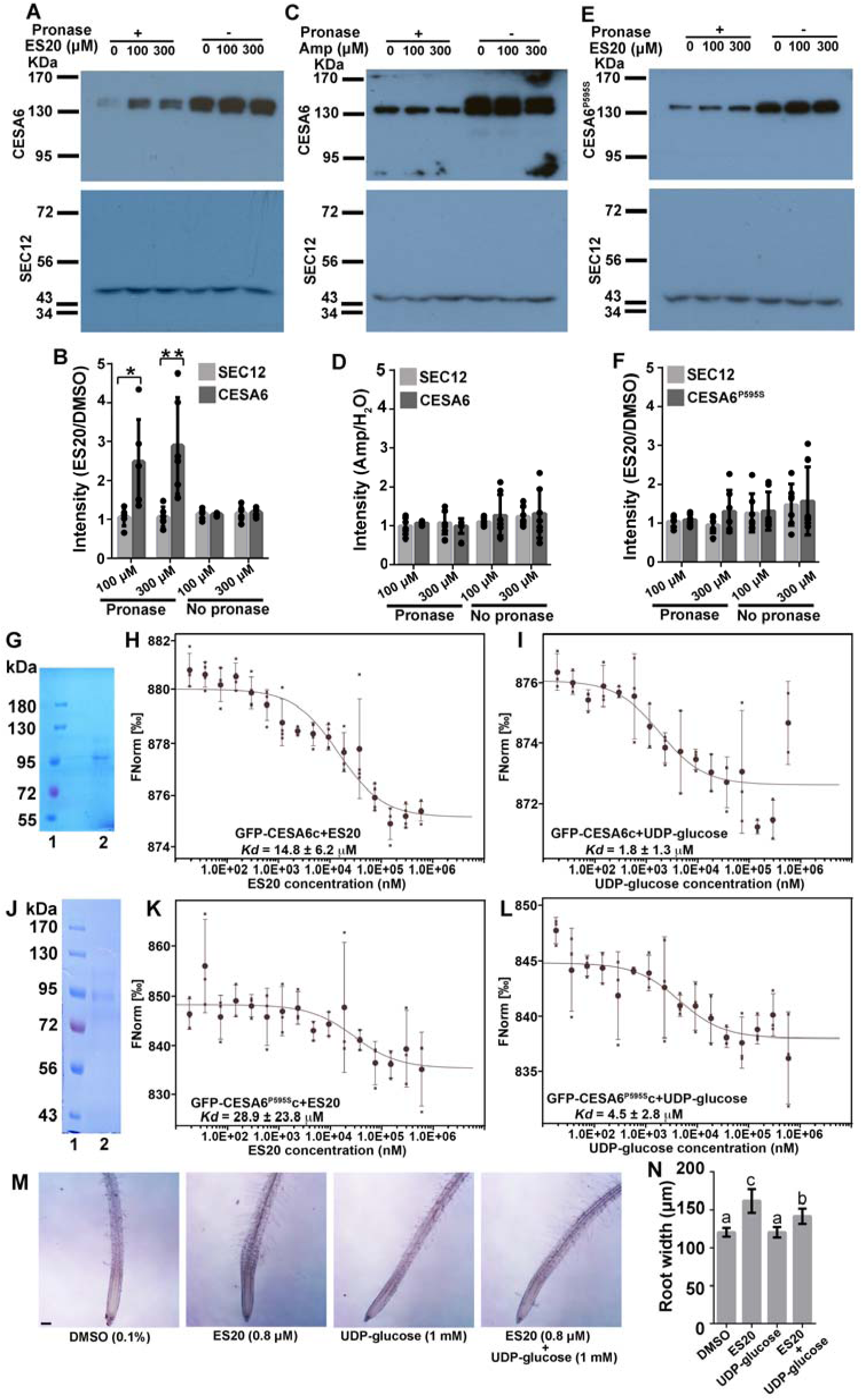
ES20 directly interacts with CESA6. **A** to **F**, ES20 interacts with CESA6 in DARTS assay. **A**, **C**, and **E**, Representative western blots of DARTS assays for YFP-CESA6 with ES20 (**A**), YFP-CESA6 with Ampicillin (**C**), and YFP-CESA6^P595S^ with ES20 (**E**), respectively. **B**, **D**, and **F**, Quantitative analysis of DARTS assays for YFP-CESA6 with ES20 (**B**), YFP-CESA6 with Ampicillin (**D**), and YFP-CESA6^P595S^ with ES20 (**F**), respectively. **G** to **I**, the central cytoplasmic domain of CESA6 interacts with ES20 and UDP-glucose in MST assay. **G**. Purified GFP-CESA6c with a His-SUMO tag (lane 2). **H**, Thermophoresis binding curve shows direct interaction between GFP-CESA6c and ES20. **I**, Thermophoresis binding curve shows direct interaction between GFP-CESA6c and UDP-glucose. **J** to **L**, the central cytoplasmic domain of CESA6^P595S^ interacts with ES20 and UDP-glucose in MST assay. **J**. Purified GFP-CESA6^P595S^c with a His-SUMO tag (lane 2). **K**, Thermophoresis binding curve shows direct interaction between GFP-CESA6^P595S^c and ES20. **L**, Thermophoresis binding curve shows direct interaction between GFP-CESA6^P595S^c and UDP-glucose. **M** and **N**, UDP-glucose could partially complement the root swollen caused by ES20. **M**, Representative images of seedlings treated with DMSO (0.1%), ES20 (0.8 μM), UDP-glucose (1 mM) and ES20 (0.8 μM) + UDP-glucose (1 mM). **N**, Quantification on the root width at the elongation zone of seedlings with different treatments as shown in **M**. The letters in **N** indicate the statistically significant differences determined by one-way ANOVA tests followed by Tukey’s multiple comparison tests in different samples. Different letters indicate significant differences between groups (p < 0.05). Error bars: mean ± SD, with *n* = 6 in **B**, **D** and **F**, *n* = 3 in **H, I, K** and **I,** and *n* = 16 in **N**. In **B**, * indicates p < 0.05 and ** indicates p < 0.01, by two-tailed student’s *t* test. Scale bar in **M**: 100 μm.

We next purified the central cytoplasmic domain of CESA6 (amino acids 322–868) with a GFP tag (GFP-CESA6c) (Figure 5G) and used a Microscale Thermophoresis (MST) assay (Wienken et al., 2010; Jerabek-Willemsen et al., 2011) to detect the interaction between ES20 and GFP-CESA6c. MST detects the movement of biomolecules as a function of ligand in the presence of a temperature gradient. The thermophoresis of a protein-ligand complex often differs from a protein alone due to binding-induced changes in size, charge and solvation energy (Wienken et al., 2010; Jerabek-Willemsen et al., 2011). We detected direct interaction between ES20 and GFP-CESA6c from the thermophoresis binding curve (Figure 5H). We also performed MST assay to detect possible interaction between GFP-CESA6c and UDP-glucose. The thermophoresis binding curve indicates direct interaction between GFP-CESA6c and UDP-glucose (Figure 5I). Similar amplitude in MST assays have been reported in studies of protein-protein interaction or protein-ligand interactions (Chen et al., 2017; Liu et al., 2017; Kosmacz et al., 2018; Yan et al., 2018; Zhai et al., 2018; Gerrits et al., 2019; Stepek et al., 2019; Warren et al., 2019). However, we did not detect interaction between ES20 and GFP or between negative control molecule Ampicillin and GFP-CESA6c (Supplemental Figure, Figure S9). We next purified the central cytoplasmic domain of CESA6 carrying P559S mutation (CESA6^P595S^c) from *E.coli* (Figure 5J) and performed MST assay to test the interaction of CESA6^P595S^c with ES20 and UDP-glucose. From the thermophoresis binding curve, CESA6^P595S^c can interact with both ES20 and UDP-glucose (Figure 5K, 5L). The results of DARTS and MST assays confirm that ES20 directly interacts with CESA6. The mutation P595S abolishes the interaction between CESA6 and ES20 in DARTS assay but recombinant CESA6^P595S^c can still interact with ES20. The results from DARTS and MST assays are probably not directly comparable because DARTS assay uses endogenous CESA6 protein that is part of the protein complex in the lipid-bilayer environment while the recombinant CESA6 cytoplasmic domain does not contain the transmembrane domain. It won’t be surprising that the cytoplasmic domain cannot fold into exact the same conformation as the endogenous full-length CESA6 in the cell.

To experimentally test whether ES20 compete with UDP-glucose for the catalytic site, we examined whether externally supplement of UDP-glucose could partially compensate the inhibitory effects of ES20. Previous studies indicate that efficient penetration of externally supplemented UDP-glucose through cell membrane often requires cell wounding (Brett, 1978; Susette and Gordon, 1983). However, more recent reports also show that externally supplemented UDP-Glucose could rescue male fertility defects of UDP-Glucose deficiency mutants and reverse the inhibitory effect of an UGPase/USPase inhibitor in pollen (Park et al., 2010; Decker et al., 2017). We co-treated wildtype seedlings with ES20 (0.8 μM) and UDP-glucose (1 mM) and used DMSO (0.1%, v/v), UDP-glucose (1 mM) and ES20 (0.8 μM) alone as control treatments. After overnight incubation, we found that externally supplemented UDP-Glucose could not completely reverse the effects of ES20, because the seedlings co-treated with 1 mM UDP-Glucose and 0.8 μM ES20 still had significant swollen roots (Figure 5M and 5N). However, we consistently detected statistically differences in root width between seedlings treated with 0.8 μM ES20 alone and seedlings co-treated with 1 mM UDP-Glucose and 0.8 μM ES20 (Figure 5M and 5N). The roots in ES20 and UDP-glucose co-treated samples had less swollen phenotypes when compared with roots treated with ES20 alone (Figure 5M and 5N). UDP-glucose alone treatment did not affect root width. Statistically significant compensation of ES20 inhibitory effect by UDP-glucose is consistent with our modeling results that ES20 targets CESA6 at the catalytic site.

### Inhibition of CESA catalytic activity by ES20 treatment affects CSC celluar localization reduces CSCs delivery to the PM

We next used ES20 as an inhibitor of plant CESA catalytic activity and investigated whether altered catalytic activity affected CSC dynamic behavior. Functional fluorescence-tagged CESAs have been found to localize mainly at the Golgi, PM, and small CESA compartments (SmaCCs) (Paredez et al., 2006; Crowell et al., 2009; Gutierrez et al., 2009; Zhang et al., 2019). At the PM, CSCs translocate along cortical microtubules with a velocity that is dependent upon catalytic activity (Paredez et al., 2006; Gutierrez et al., 2009; Fujita et al., 2013; Morgan et al., 2013). To confirm that ES20 inhibits the synthesis of β-1,4-glucan, we treated *Arabidopsis* seedlings expressing YFP-CESA6 in the *prc1-1* background with 6 µM ES20 or 0.1% DMSO for 30 min and imaged the PM of root epidermal cells with spinning disk confocal microscopy (SDCM), as described previously (Zhang et al., 2019) (Figure 6A-6C). Time projections from 5-min time-lapse series showed linear tracks in mock-treated cells, whereas ES20-treated cells had fewer tracks (Figure 6A). By analyzing kymographs from multiple cells and roots, we found that ES20 treatment significantly reduced the rate of CSC motility to 74 ± 36 nm/min compared to 137 ± 65 nm/min for mock-treated cells (Figure 6B, 6C). Reduced CSC velocity after ES20 treatment is consistent with our molecular docking results that ES20 inhibits cellulose polymerization.

**Figure 6.**
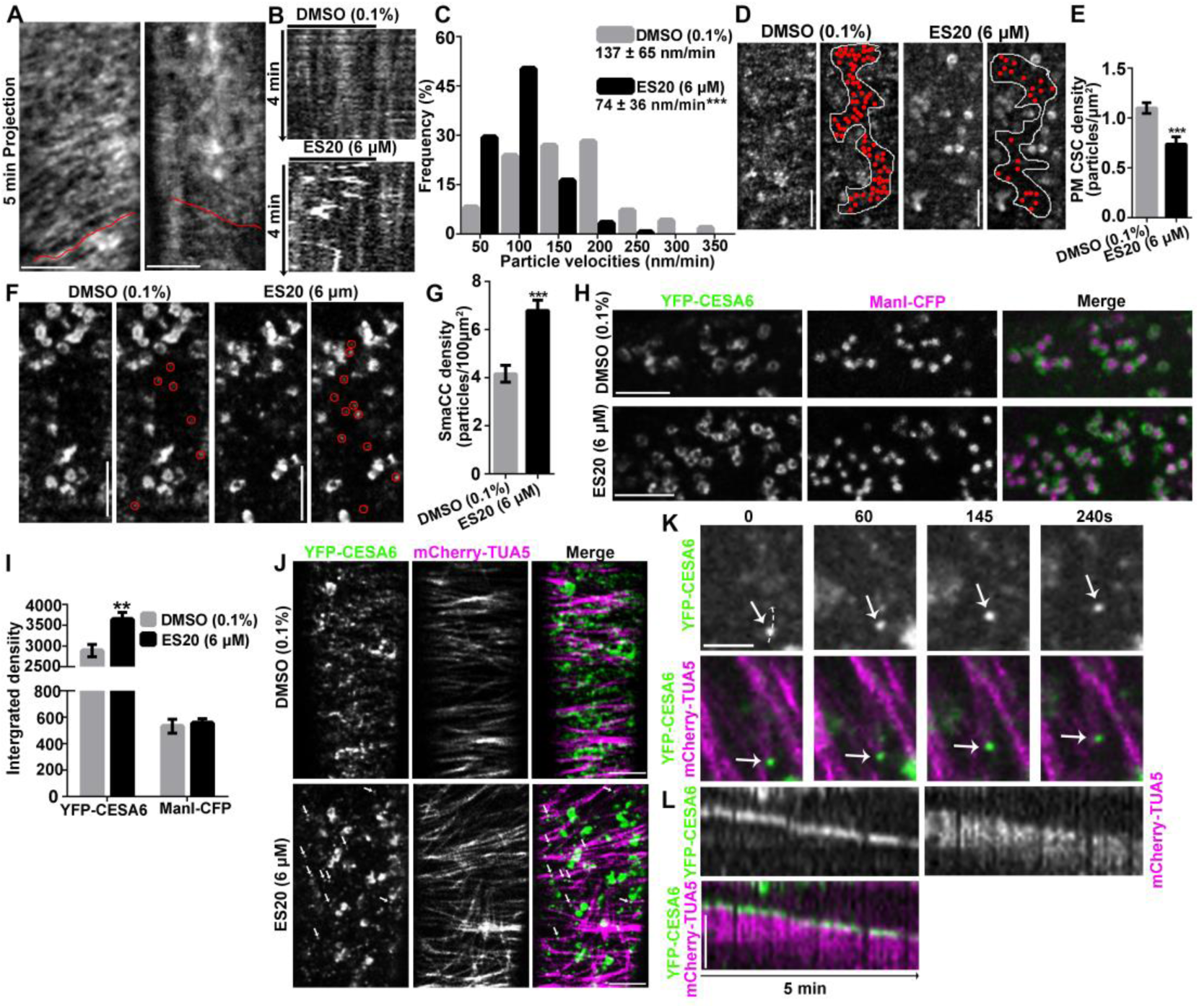
ES20 reduces CSC localization at the PM and increases CSC at the Golgi. **A** to **C**, ES20 reduced the velocity of CSCs at the PM. **A**, Representative time projections using average intensity images from a time-lapse series of YFP-CESA6 particles in root epidermal cells. **B**, Kymographs of trajectories marked in (**A**). **C**, Histogram showed the frequencies of YFP-CESA6 particle velocity after treatment with 0.1% DMSO or 6 μM ES20 for 30 min. Data represent mean ± SD (*n* = 320 CSC trajectories from 18 seedlings per treatment). **D** and **E**, ES20 reduced PM-localized YFP-CESA6 in root epidermal cells after ES20 treatment. Representative images (**D**) and quantification (**E**) of PM-localized YFP-CESA6 in root epidermal cells after 0.1% DMSO or 6 μM ES20 treatment were shown. Data represent mean ± SE (*n* = 20 cells from 10 seedlings). **F** and **G**, The density of cortical SmaCCs, as indicated by red circles, was increased by ES20 treatment (30 min). Data represent mean ± SE (*n* = 20 cells from 10 seedlings per treatment). **H** and **I**, ES20 increased the abundance of CSC at the Golgi. **H**, Representative images of Golgi-localized YFP-CESA6 and ManI-CFP after 0.1% DMSO (top) or 6 μM ES20 (bottom) treatment for 1 h. **I**, Quantification of integrated fluorescence intensity of Golgi-localized CSCs and ManI as described in (**H**). Data represent mean ± SE (*n* = 60 from 14 seedlings). **J**, CSCs were depleted from the PM after treatment with 6 μM ES20 for 2 h, whereas microtubule-associated CESA compartments accumulated, as indicated by white arrows. **K**, Magnified view on the association of CESA compartment (pointed by white arrows) with microtubules in time course image after 6 μM ES20 treatment for 2h. **L**, Kymograph image to show the association of CESA compartment with the microtubules as shown in **K**. Scale bars, 5 μm. ** indicates p < 0.01 and *** indicates p < 0.001 by two-tailed student’s *t* test.

CESA subunits are thought to be assembled into CSC rosettes at either the ER or at the Golgi, based on observation of rosettes in Golgi membranes of freeze-fractured cells examined by electron microscopy (Haigler and Brown, 1986; Gardiner et al., 2003). Golgi-localized CSCs are mainly delivered to the PM via putative secretory vesicles which may be the same as SmaCCs (Gutierrez et al., 2009; Zhang et al., 2019). To quantify trafficking and dynamics of CSCs within and between compartments, we performed both static and time-lapse analyses of YFP-CESA6 localization by collecting 3- and 4-D stacks of images from epidermal cells in the root elongation zone with SDCM. We found that after 30 min of 6 μM ES20 treatment, the density of PM-localized CSCs was reduced by about 25%, from 1.1 ± 0.1 particles/μm^2^ to 0.7 ± 0.1 particles/μm^2^ (Figure 6D, 6E). In normal growing cells, some CSCs are found to localize to motile SmaCCs in the cortical cytoplasm, and their abundance rapidly increases when secretion is inhibited (Crowell et al., 2009; Gutierrez et al., 2009; Zhang et al., 2019). The exact identity and function of these SmaCCs is not well understood, but they have been found to have partial overlap with TGN proteins and are major vesicle compartments associated with CSC delivery to the PM (Gutierrez et al., 2009; Zhang et al., 2019). We found that the abundance of cortical SmaCCs was significantly increased after 30 min of 6 μM ES20 treatment, from 4.2 ± 0.4 particles/100 μm^2^ to 6.8 ± 0.4 particles/100 μm^2^ (Figure 6F, 6G).

Live-cell imaging of functional fluorescence-tagged CESAs reveals abundant CSCs at the Golgi (Crowell et al., 2009; Gutierrez et al., 2009; Zhang et al., 2019). Further, immunogold labeling with an anti-GFP antibody in GFP-CESA3 plants shows CESAs are localized at the periphery of Golgi cisternae and the TGN (Crowell et al., 2009). We found that after 1 h of 6 μM ES20 treatment, the fluorescence intensity of YFP-CESA6 at Golgi was increased by more than 20% compared to the DMSO control, from 2890 ± 147 to 3642 ± 169 (Figure 6H, 6I). In contrast, the fluorescence intensity of another Golgi-localized protein, ManI-CFP that is expressed in the same cells, was not affected by ES20 treatment (Figure 6H, 6I). When we extended the 6 μM ES20 treatment to 2 h, PM-localized CSCs were completely depleted from the PM and we found increased abundance of CESA compartments that were associated with microtubules in the cortical cytoplasm (Figure 6J-6L, Supplemental Movie 1). These data demonstrate that ES20 treatment affects proper subcellular localization of CSCs.

Reduced CSC density at the PM and increased CESA6 abundance in a population of cortical SmaCCs following ES20 treatment suggest that ES20 affects CSC delivery to the PM. We examined the effect of ES20 on CSC delivery to the PM using a Fluorescence Recovery After Photobleaching (FRAP) assay (Gutierrez et al., 2009; Zhang et al., 2019). We mounted YFP-CESA6 seedlings grown in normal growth media directly into media supplemented with 0.1% DMSO or 6 μM ES20 and then photobleached a small region of interest on the PM. The delivery of new CSCs to the bleached region was examined using time-lapse SDCM imaging. After a 5-min acute treatment with ES20, the delivery rate of CSCs was reduced by about 30%, from 3.0 ± 0.2 particles/μm^2^/h to 2.0 ± 0.2 particles/μm^2^/h (Figure 7A-7C). Results from the FRAP assay indicate that ES20 reduces the efficiency of CSC delivery to the PM. Increased intensity at the Golgi could result from increased CSC delivery from ER to the Golgi, reduced CSC exit from the Golgi, or both. To test this, we analyzed the dynamics of YFP-CESA6/CSC transport from ER to the Golgi using FRAP analysis. We treated YFP-CESA6 seedlings with 0.1% DMSO or 6 μM ES20 for 1 h in the presence of a low concentration of Latrunculin B (2 μM) to immobilize Golgi, followed by photobleaching of individual Golgi stacks and time-lapse imaging to examine fluorescence recovery. We found that about 30% of the photobleached YFP-CESA6 fluorescence intensity could be recovered from both DMSO- and ES20-treated seedlings within 5 min, but the rate of fluorescence recovery was not significantly different between the two treatments (Figure 7D, 7E). Increased YFP-CESA6 fluorescence intensity at the Golgi and normal delivery rate from ER to the Golgi indicate that ES20 likely affects CSC transport out of Golgi but not for transport from ER to the Golgi.

**Figure 7.**
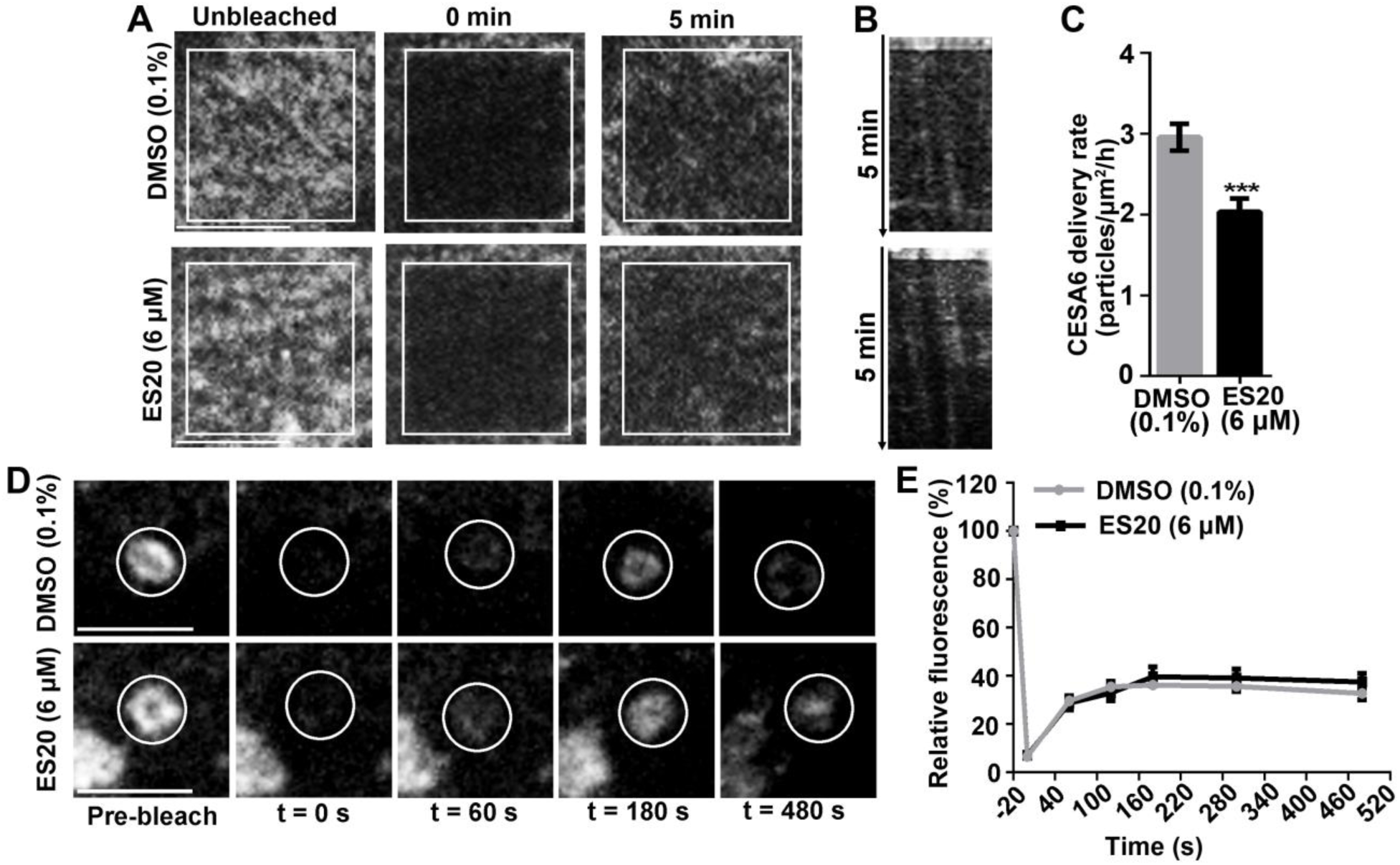
ES20 treatment inhibits CSC delivery to the PM but does not affect CSC trafficking from ER to the Golgi. **A** to **C**, ES20 reduced the delivery rate of CSCs to PM in root epidermal cells. **A**, Representative images of CSCs at the PM during FRAP analysis. **B,** Representative kymographs of trajectories of newly delivered CSCs after photobleaching. **C**. Quantification of CSC delivery rates based on FRAP assays described in (**A**). Data represent mean ± SE (*n* = 18 ROI from 15 seedlings). **D** and **E**, ES20 did not affect the delivery of CSCs from ER to the Golgi in root epidermal cells. **D**. Representative images of Golgi-localized YFP-CESA6 during a FRAP assay. **E**. Quantification of the relative recovery of CSCs at Golgi at different time points during FRAP assay. Data represent mean ± SE (*n* = 12 from 12 seedlings per treatment). Scale bars: 5 μm. *** indicates p < 0.001 by two-tailed student’s *t* test.

### Mutations in amino acids at the CESA6 catalytic site reduces CSC delivery to the PM

To further evaluate whether altering the catalytic site affects the trafficking of CSCs, we studied the trafficking dynamics of YFP-CESA6 with mutations at the predicted catalytic site in etiolated hypocotyl epidermal cells. We selected three transgenic lines that express YFP-CESA6 constructs carrying mutations at conserved amino acids in the catalytic core of bacterial CesA in *cesa6/prc1-1*. We used transgenic plants expressing wildtype YFP-CESA6 construct in *cesa6/prc1-1* as a control. Among the mutated CESA6 constructs we selected, CESA6^T783I^ and CESA6^D785N^ contain mutations at the conserved TED motif at the predicted catalytic site that has been shown to be essential for cellulose catalytic synthesis in bacteria (Morgan et al., 2016) (Figure 4B, Supplemental Figure, Figure S4). We also chose CESA6^Q823E^ because it is predicted to be in the catalytic site and form a hydrogen bond with ES20 (Figure 4B, 4C). Because these mutations are predicted to affect the catalytic activity of CSCs, it is expected that the CSCs have reduced motility at the PM. By quantifying time-lapse images of CSC translocation in the PM, we found that all three mutations cause reduced motility of CSCs at the PM compared wildtype CESA6 (Figure 8A-8C). Slower motility suggests that these mutations affect efficient cellulose catalytic synthesis on the PM. We next used FRAP analysis to test whether the mutations affect the delivery of CSCs to the PM. We found that all three mutations significantly reduced the delivery rate of CSCs to the PM (Figure 8D, 8E). Both reduced motility and delivery rate observed with CESA6^T783I^, CESA6^D785N^, and CESA6^Q823E^ mutations indicate that the catalytic site of CSCs affects their subcellular trafficking.

**Figure 8.**
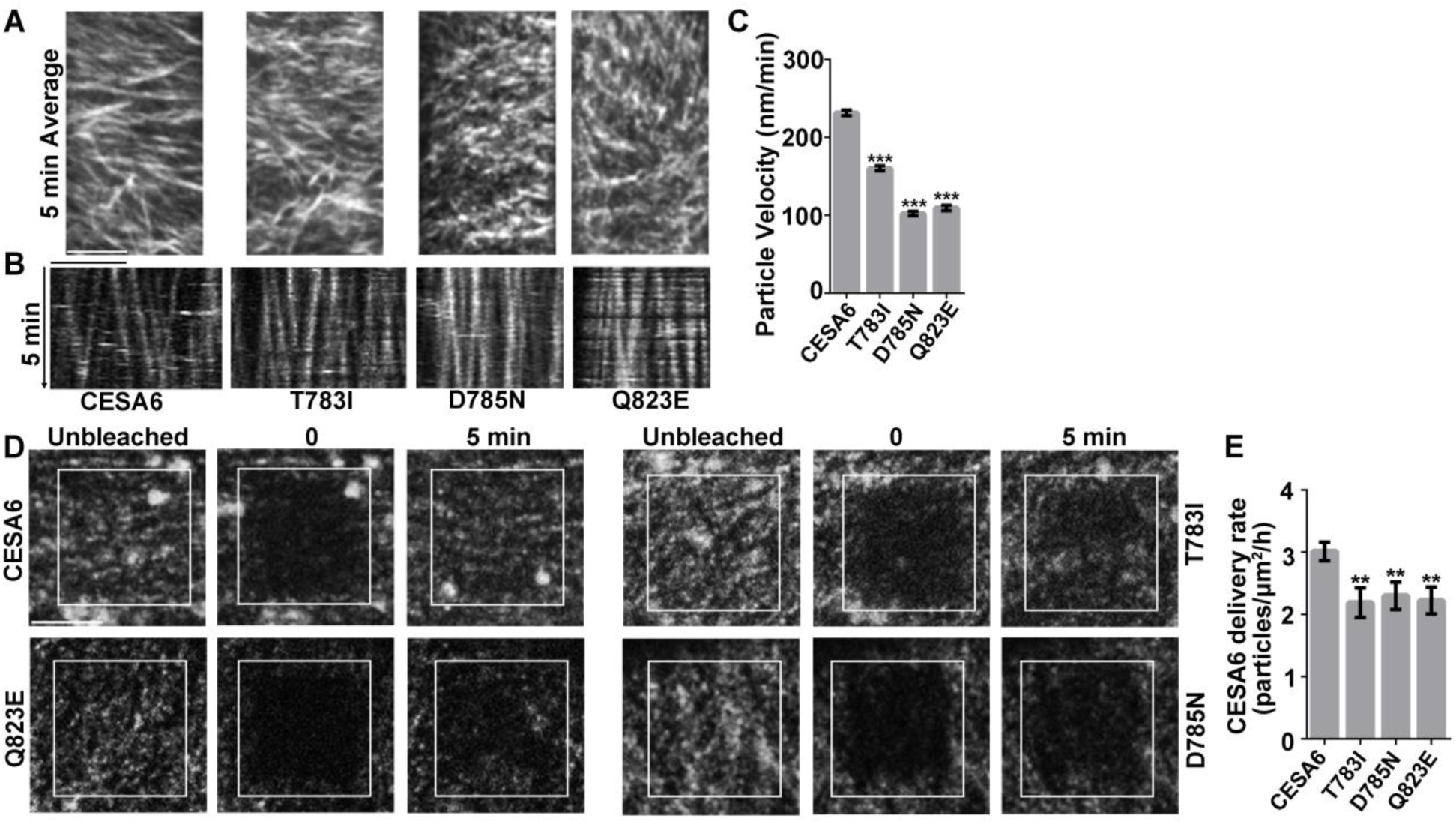
Mutations in amino acids at the catalytic site of CESA6 reduce CSC motility at the PM and reduce CSC delivery to the PM in etiolated hypocotyl cells. **A** to **C**, Mutations in amino acids at the catalytic site of CESA6 reduced the velocity of CSCs at the PM. **A**, Representative time projections of average intensity images from a time-lapse series of CSC particles from YFP-CESA6 lines carrying different mutations. For each time projection, 61 frames collected at 5-s intervals were used. **B**, Kymographs of trajectories marked in (**A**) showed the movement of CSCs over 5 min. **C**, Quantification of the velocities of CSCs at the PM in YFP-CESA6 lines carrying different mutations. Data represent mean ± SE (*n* >300 CSC trajectories from 5 seedlings for each mutated line). ***, p < 0.001 by two-tailed student’s *t* test. **D** and **E**, Mutations in amino acids at the catalytic site of CESA6 affect the delivery rate of CSCs to PM. **D**, Representative images of CSCs at the PM during FRAP analysis in YFP-CESA6 lines carrying different mutations. White boxes marked the ROI for photobleaching. **E**, Quantification on CSC delivery rates based on FRAP assays described in (**D**). Data represent mean ± SE (*n* = 20 ROIs from 10 seedlings). ** represents p < 0.01 by two-tailed student’s t test. Scale bars in **A**, **B** and **D**: 5 μm.

Because these CESA6 mutants were resistant to ES20 as assayed by seedling growth, we investigated whether the mutations cause reduced sensitivity to ES20 treatment at the cellular level as well. We first analyzed the effect of short-term ES20 treatment on the motility CSCs in the PM carrying mutations at the catalytic site of CESA6. We treated light-grown seedlings of YFP-CESA6, YFP-CESA6^T783I^, YFP-CESA6^T785N^ and YFP-CESA6^Q823E^ with 6 μM ES20 for 30 min and quantified the effect of ES20 on CSC motility at the PM, CSC density at the PM, and the abundance of cortical SmaCCs in root epidermal cells. We found that the velocity of CSCs containing mutated CESA6 was not further reduced by ES20 treatment for all three mutated YFP-CESA6 lines (Figure 9A-9C). When we investigated CSC density at the PM after ES20 treatment, we found that only in the YFP-CESA6^T783I^ line was CSC density at the PM reduced by ES20 treatment. Statistical analysis also shows that although CSC density at the PM in YFP-CESA6^T783I^ was reduced by ES20 treatment, the abundance was still higher than that of YFP-CESA6 treated with ES20 (Figure 9D, 9E). In YFP-CESA6^T785N^ and YFP-CESA6^Q823E^ lines, CSC density at the PM was not affected by ES20 treatment. We next investigated cortical SmaCC density in three mutated YFP-CESA6 lines after ES20 treatment. We found that in YFP-CESA6^T783I^, SmaCC density was increased at a level similar to that of YFP-CESA6 after ES20 treatment (Figure 9F, 9G). SmaCC density was not affected by ES20 treatment in YFP-CESA6^T785N^ and YFP-CESA6^Q823E^ lines. Reduced sensitivity to ES20 treatment for CSC motility and density in the PM, as well as cortical SmaCC density in mutated YFP-CESA6 lines indicate that the amino acids at the catalytic site are important for the inhibitory effects of ES20 at the cellular level. Our analysis of three mutated YFP-CESA6 lines also indicates that the amino acids at the catalytic site contribute differently to the response to ES20 treatment at the cellular level. CSC behavior is more sensitive to ES20 treatment in YFP-CESA6^T783I^ than YFP-CESA6^T785N^ and YFP-CESA6^Q823E^.

**Figure 9.**
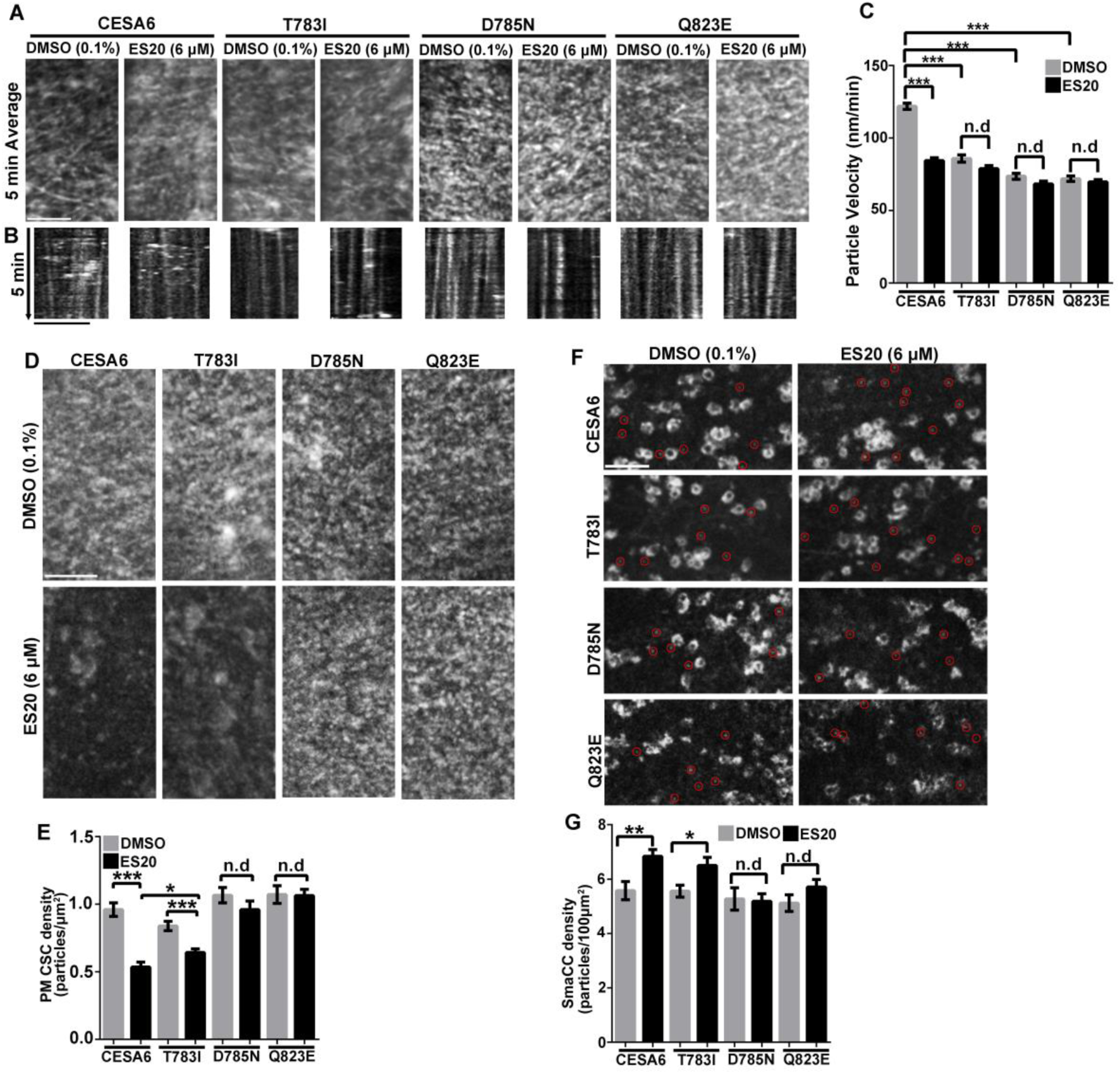
Mutations in amino acids at the catalytic site cause reduced sensitivity to ES20 treatment at the cellular level in root epidermal cells. **A** to **C**, Mutations in amino acids at the catalytic site of CESA6 reduced the sensitivity to ES20 inhibition of CSC velocity. **A**, Representative time projections of average intensity images from a time-lapse series of YFP-CESA6 particles from YFP-CESA6 lines carrying different mutations after DMSO or ES20 treatment. For each time projection, 61 frames collected at 5-s intervals were used. **B**, Kymographs of trajectories marked in (**A**) showed the movement of CSCs over 5 min. **C**, Quantification of the velocities of CSCs at the PM in YFP-CESA6 lines carrying different mutations after ES20 treatment. Data represent mean ± SE (*n* >300 CSC trajectories from 6 seedlings for each mutated line). **D** and **E**, Mutations in amino acids at the catalytic site of CESA6 cause reduced sensitivity to ES20 inhibition of CSC density at the PM. Representative images (**D**) and quantification of (**E**) of PM-localized YFP-CESA6 carrying different mutations in root epidermal cells after 0.1% DMSO or 6 μM ES20 treatment. Data represent mean ± SE (*n* = 24 cells from 12 seedlings). **F** and **G**, Some mutations in amino acids at the catalytic site of CESA6 cause reduced sensitivity to ES20 induction of cortical SmaCCs. The density of cortical SmaCCs was increased by ES20 treatment (30 min). Data represent mean ± SE (*n* = 24 cells from 12 seedlings). * represents p<0.05, ** represents p<0.01, and *** represents p < 0.001 by two-tailed student’s t test. n.d represents no significant difference. Scale bars in **A**, **D** and **F**: 5 μm.

## DISCUSSION

Due to the importance of cellulose in agriculture and industry, understanding the mechanisms of cellulose biosynthesis has been one of the most important topics in biology. Chemical inhibitors that allow transient manipulation of target protein behaviors serve as valuable tools in biological research. Here we identified a novel cellulose synthase inhibitor that targets the catalytic site of Arabidopsis CESA6. ES20 is likely to have different target site than isoxaben and C17 because the mutations that lead to reduced sensitivity to these inhibitors are very different (Scheible et al., 2001; Desprez et al., 2002; Hu et al., 2016; Hu et al., 2019). Indaziflam is another potent cellulose synthesis inhibitor but it is not known how it affects CESA activity (Brabham et al., 2014). From our mutant screen, we only identified mutants of *CESA6* but not other *CESAs* that are resistant to ES20. However, we found that comparable mutants in CESA1 (*any1*, *CESA1^D604N^*) and CESA7 (*fra5*, *CESA7^P557T^*) also have reduced sensitivity to ES20 inhibition, although a lower dosage of ES20 is required to observe a significant resistant phenotype in *any1* and *fra5*. Reduced sensitivity of *any1* and *fra5* to ES20 indicates that ES20 might target CESA1 and CEAS7, and probably other CESAs as well. We noticed that the mutations in other CESAs might have stronger growth phenotypes than that of CESA6 mutants, for example, *any1* has a stronger root growth phenotype than *es20r4* (CESA6^D605N^) (Figure 3). It is possible that ES20 can target multiple CESAs but we could not identify mutants in other CESAs because the dosage of ES20 (5 μM) we used for the screening was too high to allow us to identify those mutants. CESA7 has been shown to function mainly in secondary cell wall synthesis (Gardiner et al., 2003; Taylor et al., 2003; Brown et al., 2005). However, we found that *fra5* has a slightly reduced root growth at the young seedling stage and demonstrated a slightly reduced sensitivity to ES20. It seems that *CESA7* has a function in young seedling root development as well. We also cannot rule out the possibility that CESA6 holds a special position in the CSC rosette that allows ES20 to target only CESA6 to affect the entire protein complex during cellulose synthesis. We expect further characterization of ES20 specificity for different CESAs in Arabidopsis and other plants will be required for better use of ES20 as a CESA inhibitor. ES20 does not affect the localization of PIN2 nor BRI1, indicating it does not disrupt general exocytosis in plants. We did not find evidence for other potential targets for ES20. In our mutant screen, we did not find mutations in any other genes that have caused reduced sensitivity to ES20. Based on our current results, we expect ES20 can be used as a CESA6 inhibitor in Arabidopsis to understand the molecular mechanisms of cellulose catalytic synthesis and the integration between cellulose catalytic synthesis and CSC dynamic behaviors.

Due to the importance of cellulose in plant growth, null mutants of CESAs often have severe growth phenotypes that have limited their contributions in understanding the molecular mechanisms of CESA function. Missense mutations at critical domains can provide valuable information on the mechanisms of domain function. We obtained a group of new *cesa6* missense mutation alleles that are located at the predicted CESA6 catalytic site. These mutants have different severity in plant growth defects and have reduced sensitivity to ES20. The mutant growth defects and the reduced motility of mutated proteins on the PM support the modeled structure of CESA6 catalytic site. Site-directed mutagenesis of amino acids at the predicted catalytic site and the phenotype analyses on these newly designed mutants provide further support for our modeling results. Based on the structure modeling and molecular docking results, there are 25 amino acids located within 4 Å to UDP-glucose phosphonate and 18 amino acids located within 5 Å to ES20. Interestingly, the list of 18 amino acids that are predicted to be within 5 Å to ES20 completely overlap with the 25 amino acids that are predicted to be within 4 Å to UDP-glucose phosphonate. Among 25 amino acids that are located within 4 Å to UDP-glucose phosphonate, only L401 is located in the PCR domain (aa399-523) and none is located in the CSR domain (aa643-771), indicating the majority of the amino acids in PCR and CSR domains might not be directly involved in the catalytic process. Although further high-resolution experimental data, such as x-ray diffraction or cryo-EM, is required to resolve the structure of CSC catalytic site, our modeling results with the support of chemical genetic analysis provide important reference for understanding the catalytic site composition of plant CESAs. Future site-directed mutagenesis on other amino acids at the predicted catalytic site will allow further validation of our modeled catalytic site composition.

Years of studies have found that microtubules, actomyosin, vesicle trafficking machineries, and CESA-interacting proteins play important roles in regulating precise control of CSC trafficking and cellulose biosynthesis. Here we show, by quantitative live cell imaging of wildtype YFP-CESA6 treated with ES20 as well as mutated YFP-CESA6 at amino acids in the catalytic site, that the catalytic site of CSCs contains information that is important not only for cellulose synthesis but also for CSC delivery to the PM. Three mutations at the catalytic site, YFP-CESA6^T783I^, YFP-CESA6^T785N^ and YFP-CESA6^Q823E^, significantly reduce CSC motility on the PM and CSC delivery to the PM. It is possible that interactions with ES20 or mutation at the catalytic site alter the structural conformation of CSCs and this conformation change might lead to reduced efficiency of the CSCs being recognized by other proteins that are essential for regulating CSC delivery to the PM.

When we tested the sensitivity of mutated YFP-CESA6 to ES20 treatment at the cellular level, we found that the velocity of CSCs on the PM is not further reduced by ES20 treatment. When we compared the static localization of wildtype YFP-CESA6 and mutated YFP-CESA6 treated with ES20, we found that the mutations cause different sensitivity to ES20 treatment at the cellular level. For example, after 30 min 6 μM ES20 treatment, the density of CSCs at the PM was reduced and the density of cortical SmaCCs was increased significantly in YFP-CESA6^T783I^. However, CSC density at the PM and cortical SmaCCs density was not affected by ES20 treatment in YFP-CESA6^T785N^ and YFP-CESA6^Q823E^. Different sensitivity to ES20 treatment indicates that the amino acids at the catalytic site may play different roles in regulating CSC trafficking. We did not observe any abnormal CSC density at the PM and the abundance of cortical SmaCCs in any of the three mutated YFP-CESA6 constructs from static images (Figure 9D-9G). However, both ES20 treatment and mutations at the catalytic site significantly reduced the motility of CSC in the plane of the PM as well as the rate of CSC delivery to the PM. Our CSC motility analysis is consistent with previous finding that the catalytic activity affects the motility of CESA1^D604N^ (*any1*) at the PM (Fujita et al., 2013). The difference between ES20 treatment and catalytic site mutation in CSC PM-localized CSC density and SmaCC density may because that ES20 treatment time was short (30 min) and reflects the acute response of catalytic inhibition while analysis on mutated YFP-CESA6 reflects the equilibrated delivery and endocytic recycling that can provide feedback to each other. Thus, short-term inhibitor treatment allows a more direct observation on the cellular response of catalytic inhibition. We found that Golgi-localized CSCs were increased after 1 h ES20 treatment (Figure 6H, 6I). Due to the poor understanding on the mechanisms of CSC assembly, modification and trafficking at the Golgi, it is unclear how ES20 treatment affects CSC localization at the Golgi. It is possible that the inhibitor treatment affects the efficiency of CSC assembly or the efficiency of assembled CSC being selected by the receptors for CSC export from Golgi.

There are eight predicted transmembrane domains in each CESA. We identified two amino acids, L286 and G935, at the transmembrane domains that led to reduced sensitivity to ES20 in plant growth when they are mutated. We were not able to include these transmembrane domain amino acids in structure modeling and molecular docking and do not know how they are involved in cellulose catalytic synthesis or CESA6 interaction with ES20. During cellulose catalytic synthesis, some amino acids at the transmembrane domain must form the transmembrane pore to facilitate glucose polymerization and polymer transmembrane translocation. It is possible that amino acids L286 and G935 are involved in the transmembrane translocation of cellulose polymers. G929 is located at the small predicted cytoplasmic loop between 4^th^ and 5^th^ transmembrane helixes and the mutation at this amino acid caused the most obvious resistance to ES20. The function of this amino acid in cellulose synthesis awaits further investigation. Previous evidence has shown that the amino acids between the 5^th^ and the 6^th^ transmembrane helixes are essential for CESA function (Slabaugh et al., 2014). G935 is located at the 5^th^ transmembrane helix and G929 is very close to the 5^th^ transmembrane helix (Figure 3C). It is possible that the amino acids at the loop between 4^th^ and 5^th^ transmembrane helix and the amino acids at the 5^th^ transmembrane helix participate in cellulose catalysis and CSC relocation upon successful adding of glucose to existing cellulose chain. It is possible that the loop between the 5^th^ and 6^th^ transmembrane helixes is oriented toward the cytosol phase to provide essential structural conformation support during cellulose synthesis (Slabaugh et al., 2014).

We found discrepancy between our observation and a recent publication on the function of CESA6^Q823^ and CESA6^D395^ (Park et al., 2019). There authors created GFP-CESA6^Q823E^ and GFP-CESA6^D395N^ constructs using CESA6 coding sequence driven by endogenous promoter and tested the function of these constructs in complementing the growth phenotype of *prc1-1*. They found that GFP-CESA6^Q823E^ could completely rescue the growth phenotype of *prc1-1* but GFP-CESA6^D395N^ could not rescue at all. They also found that GFP-CESA6^Q823E^ had a similar motility on the PM compared with wildtype GFP-CESA6. In our hands, when we used the genomic content of CESA6 containing endogenous promoter and the genome sequence to create the mutation and used single insertion transformants for phenotype analysis, both YFP-CESA6^Q823E^ and YFP-CESA6^D395N^ could partially rescue the growth phenotype of *prc1-1*. The motility of YFP-CESA6^Q823E^ at the PM was significantly reduced when compared to the wildtype YFP-CESA6 (Figure 8). Different observations on the function of CESA6^Q823^ and CESA6^D395N^ might result from the differences in the constructs used for the complementation experiments. Nonetheless, the overall conclusion of both studies supports the argument that CESA catalytic activity correlates with efficient transport of CSCs through the endomembrane system.

## MATERIAL AND METHODS

### Plant materials and growth conditions

To test the inhibitory effect of ES20 on plant growth, *Arabidopsis* wildtype Col-0 plants were used. To test the effect of ES20 on cellular localization of proteins in different organelles, transgenic plants expressing fluorescence-tagged PIN2, HDEL, Got1p, VHA1-a1, ROP6, PIP2a and PGP4 were used (Cutler et al., 2000; Matsushima et al., 2003; Xu and Scheres, 2005; Dettmer et al., 2006; Cho et al., 2007; Fu et al., 2009; Geldner et al., 2009). Seeds for plants that were used for growth assays or live cell imaging were sequentially sterilized with 50% bleach and 75% ethanol. After washing with sterilized water, seeds were sowed on ½-strength Murashige and Skoog (MS) media with 1% sucrose and 0.8% agar at pH 5.8. The plants were grown under continuous light of 130 μmol m^-2^ s^-1^ intensity illuminated by Philips F25T8/TL841 25 watt bulb at 22 °C.

### Plant growth assay

To quantify the inhibitory effect of ES20 on *Arabidopsis* root growth, sterilized wildtype seeds were sowed on gridded petri plates containing ½-strength MS media supplemented with different concentrations of ES20. The plates were placed in vertical orientation in the growth chamber for root measurement. Starting from 3 d after the plates were placed in the growth chamber, the plates were scanned using Epson Perfection V550 scanner every day. The root length of plants was measured using ImageJ. To test the effect of ES20 on etiolated hypocotyl growth, sterilized wildtype seeds were sowed on ½-strength MS media supplemented with different concentrations of ES20. The petri dishes were wrapped in two layers of aluminum foil and kept at 22 °C for 7 d. The petri dishes were scanned and the hypocotyl length was measured using ImageJ. ES20 was dissolved in DMSO to obtain a stock solution of 12 mM and stored at −20 °C.

To analyze the effect of ES20 treatment on epidermal cell growth from light-grown roots, 5-day-old wildtype seedlings were treated with 0.1% DMSO or 6 μM ES20 for 12 h. The seedlings were stained with 1 μM fluorescein diacetate (ACROS organics) for 5 min and the fluorescence in epidermal cells was imaged with a Zeiss 710 laser scanning confocal microscope equipped with a 20X objective. To analyze the effects of ES20 treatment on hypocotyl cell growth, 5-day-old wildtype seedlings grown in the dark were stained with 1 μM fluorescein diacetate for 5 min and the fluorescence of epidermal cells from the middle section of the hypocotyl were imaged under the same condition as for root epidermal cells.

### EMS mutagenesis and mutant screening

In order to obtain a mutagenized *Arabidopsis* population, SYP61-CFP and PIN2-GFP seeds were mutagenized following a published protocol (Kim et al., 2006). Mutagenized seeds were sowed in soil and the plants were grown under continuous light and allowed to self, yielding M2 seeds. The M2 seeds were collected as pooled populations. About 400,000 seeds from the M2 generation of the SYP61-CFP population and 100,000 seeds from the PIN2-GFP M2 population were sterilized and sowed on media containing 5 μM ES20. Individual plants with elongated roots and green leaves were transferred to soil to produce the M3 generation. The M3 plants were examined for sensitivity to ES20. Individual M3 lines with reduced sensitivity to ES20 were crossed to *Ler* ecotype to generate the mapping population and were also crossed to SYP61-CFP or PIN2-GFP to clean up the genetic background.

### High-throughput genome sequencing and sequence analysis

The seeds from F2 populations of mutants crossed with *Ler* were sowed on media containing 5 μM ES20 and the segregation of resistant seedlings was evaluated. The F2 populations of the outcrosses segregated for sensitivity to ES20. For each mutant, about 100 F2 seedlings with longer roots on 5 μM ES20 were pooled for DNA isolation. The genomic DNA was applied for high-throughput sequencing. The resulting DNA sequence was aligned to the *Arabidopsis* genome (TAIR10) and the single nucleotide polymorphism (SNP) was analyzed. Candidate SNPs for ES20 sensitivity were identified through next-generation EMS mutation mapping tool (Austin et al., 2011).

### Crystalline cellulose content measurement

*Arabidopsis* wildtype seeds were sowed on media supplemented with 0.1% DMSO or different concentrations of ES20. After stratification, the plants were grown in the dark for 7 d or under light for 10 d. 7-day-old dark-grown seedlings or roots from 10-day-old light-grown seedlings were used for cell wall preparation. Dark-grown seedlings were washed with ddH_2_O three times to remove seed coats and any residue from the growth media, then ground into fine powder under liquid nitrogen. The roots from light-grown seedlings were cut and washed with ddH_2_O to remove any residue from the growth media, then ground into fine powder under liquid nitrogen. The powder was extracted 2 times with 80% ethanol, once with 100% ethanol, once with 1:1 (v/v) MeOH-CHCl_3_, and once with acetone. The resulting insoluble cell wall fraction was dried in a fume hood for 2 d before weight measurement. Cellulose content was measured by the Updegraff method (Updegraff, 1969; Foster et al., 2010). Briefly, cell wall material was hydrolyzed by trifluoroacetic acid (TFA) and then Updegraff reagent (acetic acid: nitric acid: water, 8:1:2 v/v) to yield crystalline cellulose. Crystalline cellulose was hydrolyzed by 72% sulfuric acid to glucose. Glucose concentration was measured with a colorimetric method by developing color in Anthrone reagent (freshly prepared 2 mg/mL anthrone in concentrated sulfuric acid) and reading OD625 nm in a plate reader (Tecan Infinite 200Pro). Nine repeats were performed for each treatment, including 3 repeats for cell wall preparation and 3 repeats for measurement.

### CESA6c protein expression and purification

To obtain CESA6 central cytosolic domain protein for MST assay, we inserted GFP coding sequence into pRSF-Duet-1 vector using *SacI* and *PstI* restriction sites. GFP coding sequence was amplified from pUBN-GFP-DEST vector. Central cytosolic domain of CESA6 (CESA6c) was amplified from *Col-0* cDNA into the C-terminal of GFP. Primers used for cloning were listed in Supplemental Table 1. Verified recombinant clone was transformed into BL21 (DE3) competent cell for protein expression. The cells were cultured and grown at 37 °C in LB media till OD600 reached 0.6. Protein expression was induced by 0.1 mM isopropyl β-D-1-thiogalactopyranoside (IPTG) at 16 °C for overnight. After overnight induction, the cells were lysed by sonication and the fusion protein was purified using HisTrap HP histidine-tagged protein purification column of AKTA pure FPLC system (GE Healthcare, Pittsburgh, PA). The purified protein was further dialyzed overnight and further purified with HiLoad 16/600 Superdex 200 pg column (GE Healthcare, Pittsburgh, PA) using AKTA pure FPLC system. Purified GFP-CESA6c protein was further identified by SDS/PAGE. CESA6c construct was used as a template for creating CESA6^P595S^c construct by site-directed mutagenesis. CESA6^P595S^c protein was purified using the same protocol as CESA6c except that enriched TB media was used to grow *E.coli* cells.

### MST assays

MST assays were carried out using a Monolith NT.115 (NanoTemper) machine at the Chemical Genomics Facility of Purdue University. Increasing concentrations of ES20 were titrated against 100 nM of the GFP-CESA6c protein in a standard MST buffer (50 mM Tris, pH 7.5, 150 mM NaCl, 10 mM MgCl_2_, 0.05% Tween 20). ES20 was dissolved in DMSO and the final concentration of DMSO was 5% (vol/vol). MST standard capillaries were used to load the samples to the MST instrument. Triplicate reactions were performed for each test. The MST data was processed using MO. Affinity Analysis Version 2.3 software.

### DARTS assays

To test for the interaction between CESA6 and ES20 using DARTS assay, 7 days old YFP-CESA6 light grown seedlings were harvested and ground to powder in liquid nitrogen. The ground tissue was homogenized in the lysis buffer (50 mM Tris-Hcl, PH7.5, 150 mM NaCl, 0.5% Triton X-100, 2 mM DTT, one tablet/50 mL EDTA free Pierce protease inhibitor (Thermo Fisher)) at 2:1 ratio (2 mL buffer: 1 g tissue). Homogenized samples were transferred to a 2 mL microcentrifuge tube and centrifuged for 30 min (20,000 *g*, 4°C). The supernatant was collected after centrifugation and saved as total extracted protein. 700 μL extracted total protein was incubated with DMSO (0.1%) or ES20 (300 μM) at room temperature on an orbital shaker for 1 h. Then the mixture was divided into 6 small tubes with each contained 100 μL of the mixture and was incubated with 1 μL of pronase at 1:300 dilutions at room temperature for 30 min. The proteolysis reaction was terminated by adding SDS loading buffer and boiled at 100 °C for 6 min. The boiled samples were loaded to SDS/PAGE for further Western blot analysis. YFP-CESA6 protein was detected using anti-GFP antibody (Takara, catalog # 632381) and SEC12 was detected using anti-SEC12 antibody (Bar-Peled and Raikhel, 1997) as a control. Horseradish peroxidase conjugated secondary antibodies and Clarity Western ECL substrate (BIO-RAD, Hercules, CA) were further used to detect the presence of YFP-CESA6 and SEC12. The X-ray films were scanned and the signal intensity of each protein band was quantified after background subtraction using Image J. The relative intensities were quantified by dividing ES20 treated samples by DMSO treated samples for each pronase concentration.

### Lignin and callose staining

To examine the effect of ES20 on lignin and callose accumulation, *Col-0* seedlings were grown on 1/2 MS media supplemented with DMSO (0.1%) or ES20 (4 μM) in dark condition for three days. Lignin staining was performed following published protocol (Pradhan Mitra and Loque, 2014). Dark-grown seedlings were incubated in phloroglucinol (Acros Organic) solution (20 mg/mL in ethanol: hydrocholoric acid (2:1 vol/vol)) for 5 min and then imaged under white light. Callose staining was performed by following published protocol (Harris et al., 2012). Dark-grown seedlings were incubated in aniline blue (Acros Organic) staining solution (0.1 mg/mL in 0.07 M sodium phosphate buffer, pH 9) in the dark for 20 min. The seedlings were then imaged under UV light.

### UDP-glucose complementation on the effect of ES20 in causing root swollen

To test whether supplement of UDP-glucose can complement the effect of ES20 treatment, 3.5 days old Col-0 seedlings grown on ½ MS agar media under light condition were used. For each treatment, 16 seedings were transferred from ½ MS agar plate to 2 ml ½ MS liquid media supplemented with DMSO (0.1%, v/v), ES20 (0.8 μM), UDP-glucose (1 mM) or ES20 (0.8 μM) and UDPG (1 mM) in 24 well plate. After 17 hours of treatment, seedlings were mounted between two stripes of double-sided tape on glass slide and covered with coverslip carefully for image collection under white light using a compound microscope. The width of root elongation zone for each seedling was quantified by ImageJ.

### Vector construction and generation of transgenic Arabidopsis plants

To construct a YFP-CESA6 binary vector, a 2245bp CESA6 promoter fragment was amplified with CESA6P-F TCTGATCCAAGCTCAAGCTAAGCTTTTTCTATTCTATAGTCTTGAAAATT and CESA6P-R ATTTGTCTGAAAACAGACACAG primers using Col-0 genomic DNA as a template. The *YFP* tag was amplified from pUBN-YFP-Dest plasmid with primers YFP-F TGTCTGTTTTCAGACAAATATGGTGAGCAAGGGCGAGG and YFP-R CGACCACCGGTGTTCATCTTGTACAGCTCGTCCATG. *CESA6* with terminator was amplified from Col-0 genomic DNA with primers CESA6g-F ATGAACACCGGTGGTCGGTT and CESA6g-R GGTACCCGGGGATCCTCTAGAGTGATCCACATCTTAAATATATTA. pH7WGR2 plasmid was digested with *Hind*III and *Xba*I to remove the 35S promoter and the RFP tag. The modified pH7WGR2 linear vector without 35S promoter and RFP tag was ligated with *CESA6* promoter, *YFP* and *CESA6* genomic sequence through Gibson Assembly method using a Gibson Assembly Master Mix kit (New England Biolabs, Ipswich, MA). The construct was verified by DNA sequencing. All mutated YFP-CESA6 constructs used the verified YFP-CESA6 plasmid as the template and were obtained by Q5 Site Directed Mutagenesis Kit (New England Biolabs, Ipswich, MA) with primers listed in Supplement Table1. All of the mutated YFP-CESA6 constructs were verified by DNA sequencing. Verified constructs were further dipped into *prc1-1* (CS297) which was obtained from the Arabidopsis Biological Resource Center (ABRC) using *Agrobacterium tumefaciens* mediated transformation (Clough and Bent, 1998).

### Live-cell imaging of fluorescence-tagged proteins

To test the effect of ES20 on cellular localization of fluorescence-tagged proteins, transgenic plants expressing different fluorescence-tagged proteins were grown on ½-strength MS agar plates for 5 d. The seedlings were incubated in ½-strength MS liquid media supplemented with 6 μM ES20 for 2 h. The images were collected using a Zeiss 710 laser scanning confocal microscope equipped with a 40X/1.2 NA water objective. For imaging GFP-tagged proteins, the 488-nm laser line was used as excitation source and emission light at 493–598 nm was collected. For imaging YFP-tagged proteins, the 514-nm laser line was used as excitation source and the emission light at 519–621 nm was collected. To characterize the cellular localization and trafficking dynamics of different mutated CESA6 lines, 3 days old transgenic plants expressing YFP-CESA6 with different mutations were grown in dark on 1/2 MS (without sucrose). The 7th or the 8th cell below the hook (about 2 mm below the hook) was used for image collection.

### Structure modeling of CESA6 cytoplasmic domain

The general topology of CESA6 used in Figure 3C was predicted by TMHMM Server v. 2.0 program provided by Department of Bio and Health informatics at Technical University of Denmark (http://www.cbs.dtu.dk/services/TMHMM/). The large cytoplasmic domain of *Arabidopsis* CESA6 protein sequence (amino acids 322–868) was sent to i-TASSER server for 3D structure modeling with threading method (Roy et al., 2010; Yang et al., 2015). The modeled structure was visualized using the PyMol software (Alexander et al., 2011). The binding site of UDP-glucose on CESA6 large cytoplasmic domain model was predicted by COACH server and UDP-glucose phosphonate structure was used for the prediction per program suggestion (Yang et al., 2013b, a). The small molecule ES20 was docked with CESA6 large cytoplasmic domain model using Autodock Vina in PyRx software (Trott and Olson, 2010; Dallakyan and Olson, 2015).

### Spinning-disk confocal microscopy (SDCM)

For SDCM live cell imaging, seedlings were grown vertically for 5 d and images were taken from the 2nd or 3rd epidermal cell below the first obvious root hair initiation in the root elongation zone. Two thin strips of double-sided adhesive tape were placed on top of glass slides about 2 cm apart. 100 μl of ½-strength MS liquid growth media containing DMSO or specified concentrations of ES20 was applied to the slide and seedlings were mounted in the liquid media. A 22 x 40 mm cover glass was placed on top of the double-sided tape for imaging. For longer term imaging during CESA velocity analyses, seedlings were mounted on a piece of 1-mm thick 0.6% phytagel pad affixed to the glass slide to minimize compression and liquid evaporation.

To examine the cellular localization of YFP-CESA6; ManI-CFP and YFP-CESA6;mCherry-TUA5, SDCM imaging was performed using a CSU-X1-A1 Yokogawa scanning unit mounted on an Olympus IX-83 microscope, equipped with a 100X/1.4NA UPlanSApo oil objective (Olympus) and an Andor iXon Ultra 897BV EMCCD camera (Andor Technology). YFP, CFP and mCherry fluorescence were excited with 515-nm, 445-nm and 561-nm laser lines and emission collected through 542/27-nm, 479/40-nm and 607/36-nm filters, respectively.

For fluorescence recovery after photobleaching (FRAP) experiments, images were collected using a Zeiss Observer Z.1 microscope, equipped with a Yokogawa CSU-X1 head and a 100X/1.46 NA PlanApo objective (Zeiss). For the PM-localized CESA6 FRAP, photobleaching was performed with a Vector scanner (Intelligent Imaging Innovations) with a 515-nm laser line at 100% power and 1 ms/scan. Timelapse images were collected at the PM with a 5-s interval for 121 frames, with photobleaching in a small region (44.2 µm^2^) after the 4th frame, and recovery for total 10 min. For FRAP of CESA6-containing Golgi, a 515-nm laser line was set to 100% power with 3 ms/scan. Timelapse images were collected at the cortical cytoplasm (about 0.4 µm below the PM) with 5-s intervals for 121 frames. Photobleaching of a small region (7.1 µm^2^) was performed after the 4th frame, and recovery measured for 10 min.

### SDCM image processing and quantification

Image analysis was performed using Fiji/ImageJ (Schindelin et al., 2012). For CESA particle density analyses, ROIs that avoid abundant Golgi signals were chosen using the Freehand selection tool. CESA particles were detected automatically on 8-bit images using the Find Maxima tool with the same noise threshold for all images. CESA particle density for each ROI was calculated by dividing the number of particles by the ROI area. For CESA particle dynamic analyses, 5-min timelapse series with 5-s intervals were collected. Average intensity projections were generated to identify the trajectories of CSC particles. Image drift was corrected by the StackReg plugin (Thevenaz et al., 1998). Kymographs were generated and velocities of CESA particles were measured as the reciprocal of the slope of individual CESA particles in kymographs. For the quantification of cortical vesicles, 1 μm z-series stack with 0.2 μm as step size and 20-s timelapses were collected. Focal plane at 0.4 μm below the PM was used for the cortical SmaCC analyses. The small particles show motility in timelapse series were considered as the SmaCCs. For the FRAP assay of PM-localized CSCs, a smaller area (16 µm^2^) within the bleached region was used for analyses. The CSC delivery events during the first 5 min of recovery were manually counted according to the criteria described previously (Li et al., 2016). The particles that exhibited steady linear movement at the PM were considered as new delivery events. The CSC delivery rate was calculated by dividing the number of delivery events by the measured area and time. For the FRAP assay of CESA-containing Golgi, an area (7.1 µm^2^) within the bleached region was used for analyses. To measure the fluorescence intensity, the integrated fluorescence at selected region at different time points was calculated by subtracting the background fluorescence outside of the cell with the same size of area. The relative fluorescence of different time points was calculated by dividing the integrated fluorescence of different time points by integrated fluorescence before photobleaching. For Golgi-localized YFP-CESA6 and YFP-CESA6;ManI-CFP fluorescence analyses, 10-s timelapse series with 2-s intervals, and 1 μm z-series stack with 0.2 μm step size, were collected. Single Golgi which did not cluster with other Golgi and the z position with the highest fluorescence intensity and largest diameter was selected for analyses. A square box with an area of 3.5 µm^2^ was drawn to include the whole Golgi for analyses. The integrated fluorescence was calculated by subtracting the background fluorescence outside of the cell with the same size of area.

### Use western blot to detect the abundance of CESA6 in transgenic lines in the absence and presence of ES20

To quantify CESA6 protein level in different transgenic lines expressing mutated YFP-CESA6 in *prc1-1*, total proteins isolated from 6 days old light grown seedlings of different transgenic lines were used. About 50 seedlings from each transgenic line were treated with DMSO (0.1%) or ES20 (6 μM) for 2 hours in liquid ½ MS medium. After treatment, the seedlings were grounded into fine powder with liquid nitrogen and homogenized with lysis buffer (50 mM Tris-Hcl, PH7.5, 150 mM NaCl, 0.5% Triton X-100, 2 mM DTT) with EDTA free protease inhibitor (Thermo Fisher) at 1:1 ratio (1 mL buffer: 1 g tissue). Homogenized samples were transferred to a 1.5 mL microcentrifuge tube and centrifuged for 30 min at 20,000 g, 4°C. The supernatant was collected after centrifugation and saved as total protein extract.

Isolated total protein was loaded to SDS-PAGE for western blot analysis. Anti-GFP and anti-SEC12 antibodies were used to detect CESA6 and SEC12. The western blot results were detected using x-ray film. The western blot film was converted to electronic format by scanning the film into images using a scanner (<u>Epson Perfection V550</u>). The image file from each x-ray film was inverted using imageJ. To measure the intensity of each protein band, a rectangle box was drawn around each band and the integrated intensity in each box was measured using imageJ. The rectangle box in the same size as protein band was used to measure the integrated intensity in the area of background. Then the background intensity was subtracted from each protein band to obtain the real intensity of each protein band. After the real intensity for each protein band is obtained, the CESA6 abundance is normalized against SEC12 abundance for each protein sample by calculating the ratio of Intensity CESA6/Intensity SEC12 (R1). Then, the R1 value of each lane was normalized against wildtype YFP-CESA6 DMSO sample to obtain R2 for each sample. For example, R2^L365F,DMSO^ = R1^L365F,DMSO^/ R1^CESA6,DMSO^. The R2 values were analyzed using ANOVA to detect any difference in CESA6 abundance among different samples.

### ES20 synthesis

General Methods: NMR spectra were recorded on Bruker spectrometers (^1^H at 500 MHz; ^13^C at 125 MHz. Chemical shifts (δ) were given in ppm with reference to solvent signals [^1^H NMR: CDCl_3_ (7.26); ^13^C NMR: CDCl_3_ (77.2)]. All reactions were conducted under argon atmosphere and all solvents and reagents were used as obtained from commercial sources without further purification.

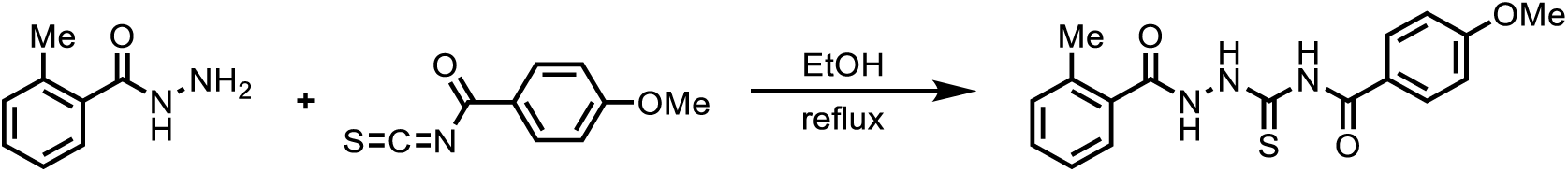

o-Toluic hydrazide (1.50 g, 10.0 mmol) was added to a stirred solution of ethanol (40 mL) followed by 4-methxoxybenzoyl isothiocyanate (1.93 g, 10.0 mmol) at room temperature. The solution was heated up to reflux under argon for 15 mins. Ethanol was removed under vacuum to give the crude product as a yellow solid. The crude product was recrystallized in ethanol to give 2.00 g purified product as a white crystal in 59% yield.

HRMS (ESI) [M + H^+^] calculated for C_17_H_17_N_3_O_3_S: 344.1063, found: 344.1064; FTIR (neat, cm^-1^) *ν*_max_ 3216, 1667, 1603, 1525, 1498, 1428, 1258, 1174, 1028, 842, 758;

^1^H NMR (500 MHz, CDCl_3_) δ: 9.48 (s, 1H), 8.95 (s, 1H), 7.87 (d, *J* = 8.8 Hz, 2H), 7.60 (d, *J* = 7.6 Hz, 1H), 7.42 (t, *J* = 7.5 Hz, 1H), 7.30 (d, *J* = 7.8 Hz, 2H), 7.01 (d, *J* = 8.9 Hz, 2H), 3.90 (s, 3H), 2.56 (s, 3H);

^13^C NMR (126 MHz, CDCl_3_) δ: 171.6, 166.0, 164.4, 164.1, 137.9, 131.6, 131.4, 129.8, 127.6, 126.0, 123.0, 114.5, 55.7, 20.2.

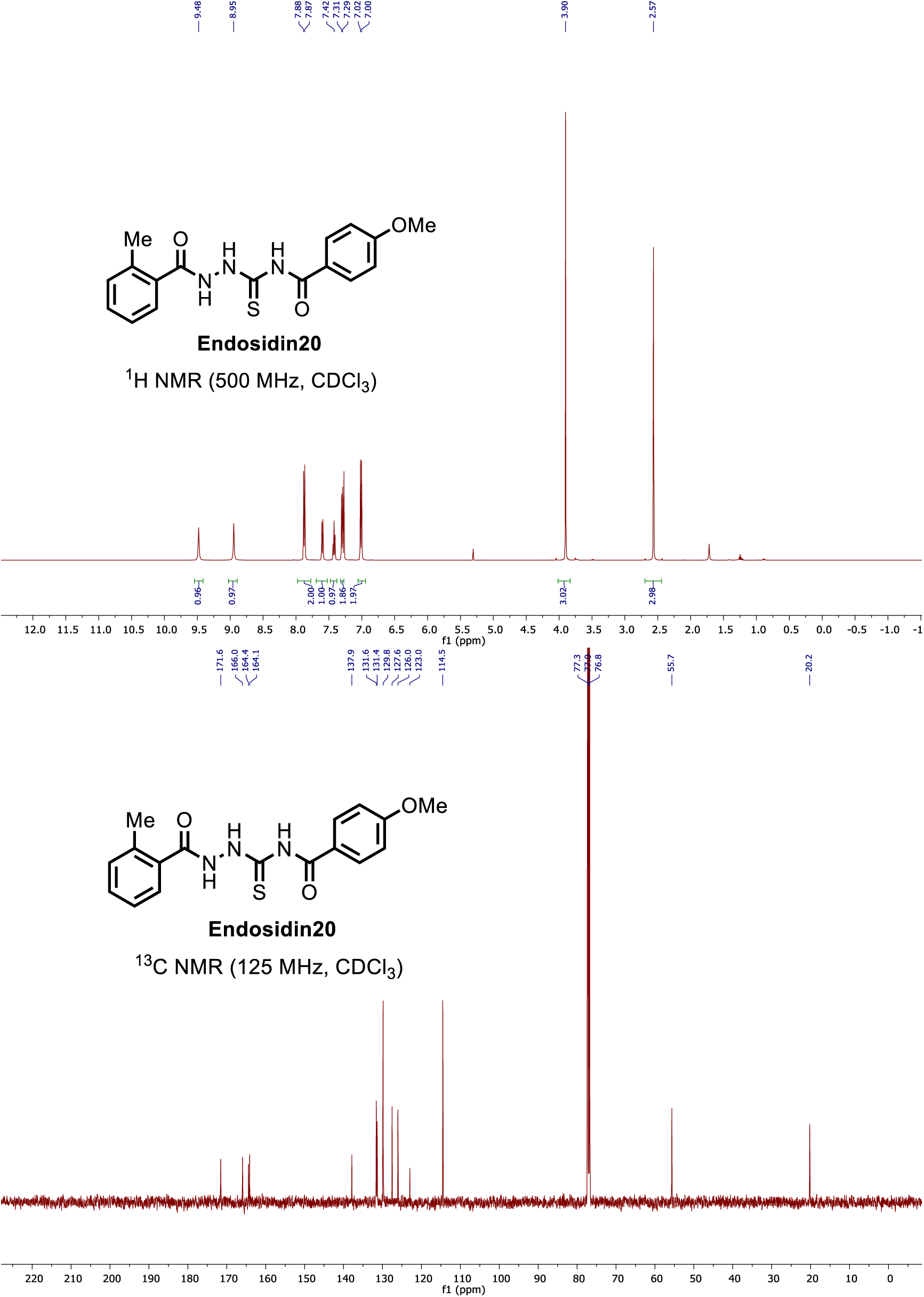

### Accession numbers

AtCESA1, AT4G32410; AtCESA6, AT5G64740; AtCESA7, AT5G17420.

## ACKNOWLEDGEMENTS

We thank Zheng-Hua Ye (University of Georgia) and Geoff Wasteneys (University of British Columbia) for providing *fra5* and *any1* seeds, respectively. We thank Ying Gu (Penn State University) for providing the YFP-CESA6;mCherry-TUA5 line. We acknowledge the Purdue Genomics facility for assistance in DNA sequencing. We are grateful to Daniel Szymanski for sharing the spinning disc confocal microscope for FRAP studies and providing us the YFP-CESA6;ManI-CFP seeds. We thank the Chemical Genomics Facility at the Purdue Institute for Drug Discovery for providing us access to the MST equipment. We thank Yun Zhou and Tesfaye Mengiste (Purdue University) for sharing the compound microscopes. Work in the Staiger laboratory on imaging of YFP-CESA6 trafficking was supported by an award from the Office of Science at the US Department of Energy, Physical Biosciences Program, under contract number DE-FG02-09ER15526. Research in Zhang laboratory was supported by Purdue University Provost’s start-up to C. Zhang.

## AUTHOR CONTRIBUTION

Lei Huang performed the mutant screening, mutant phenotype characterization, genetic complementation, CESA6 imaging and image analysis, DARTS assay, MST assay and prepared the figures. Xiaohui Li performed cell wall analysis, structure modeling, and molecular docking, and MST assay. Weiwei Zhang provided critical guidance for CESA6 imaging and image analysis. Nolan Ung performed initial compound screen and initial mutant screen. Nana Liu assisted with the biochemical assays for testing the interaction between ES20 and CESA6. Xianglin Yin and Yong Li synthesized ES20 compound. Robert Mcewan analyzed the initial high-throughput sequencing data to clone the first mutant gene. Brian Dilkes provided guidance for high-throughput sequence analysis. Mingji Dai provided guidance for ES20 synthesis. Glenn Hicks and Natasha Raikhel provided guidance for initial compound screen. Christopher Staiger provided critical guidance for CESA6 imaging and image analysis. Chunhua Zhang designed the research and wrote the article.

